# Geranylgeranylated-SCF^FBXO10^ Regulates Selective Outer Mitochondrial Membrane Proteostasis and Function

**DOI:** 10.1101/2024.04.16.589745

**Authors:** Sameer Ahmed Bhat, Zahra Vasi, Liping Jiang, Shruthi Selvaraj, Rachel Ferguson, Anish Gudur, Hagar Ismail, Ritika Adhikari, Avantika Dhabaria, Beatrix Ueberheide, Shafi Kuchay

## Abstract

E3-ubiquitin ligases (E3s) are main components of the ubiquitin-proteasome system (UPS), as they determine substrate specificity in response to internal and external cues to regulate protein homeostasis. However, the regulation of membrane protein ubiquitination by E3s within distinct cell membrane compartments or organelles is not well understood. We show that FBXO10, the interchangeable component of the SKP1/CUL1/F-box ubiquitin ligase complex (SCF-E3), undergoes lipid-modification with geranylgeranyl isoprenoid at Cysteine953 (C953), facilitating its dynamic trafficking to the outer mitochondrial membrane (OMM). FBXO10 polypeptide does not contain a canonical mitochondrial targeting sequence (MTS); instead, its geranylgeranylation at C953 and the interaction with two cytosolic factors, PDE6δ (a prenyl group-binding protein), and HSP90 (a mitochondrial chaperone) orchestrate specific OMM targeting of prenyl-FBXO10 across diverse membrane compartments. The geranylgeranylation-deficient FBXO10(C953S) mutant redistributes away from the OMM, leading to impaired mitochondrial ATP production, decreased mitochondrial membrane potential, and increased mitochondrial fragmentation. Phosphoglycerate mutase 5 (PGAM5) was identified as a potential substrate of FBXO10 at the OMM using comparative quantitative mass spectrometry analyses of enriched mitochondria (LFQ-MS/MS), leveraging the redistribution of FBXO10(C953S). FBXO10, but not FBXO10(C953S), promoted polyubiquitylation and degradation of PGAM5. Examination of the role of this pathway in a physiological context revealed that the loss of FBXO10 or expression of prenylation-deficient-FBXO10(C953S) inhibited PGAM5 degradation, disrupted mitochondrial homeostasis, and impaired myogenic differentiation of human iPSCs and murine myoblasts. Our studies identify a mechanism for selective E3-ligase mediated regulation of mitochondrial membrane proteostasis and metabolic health, potentially amenable to therapeutic intervention.

## Introduction

The ubiquitin proteasome system (UPS) controls protein turnover throughout the cell, including the cytoplasm, nucleoplasm and diverse membrane compartments ^1,2^. However, the operation of the UPS within specific membrane compartments, such as mitochondria, endoplasmic reticulum (ER), plasma membranes (PM), and other subcellular membrane compartments is not well understood. Within this framework, approximately 600 predicted E3 ubiquitin ligases (E3s), as encoded in the human genome, serve as principal nodes for substrate specificity by directly recruiting protein targets in response to internal and external cues for the UPS mediated protein turnover^1,3,4^. Although various E3s operating in cytosolic and nuclear compartments have well-established mechanisms for substrate recruitment, the membrane compartment-specific substrate recruitment and turnover by E3s is not completely clear. In particular, the full complement of E3s operating from membrane compartments and mechanism(s) by which E3s are targeted to specific membrane compartments to enable ubiquitylation of organellar substrates is not established. The processes by which E3s, especially non-transmembrane E3s, are delivered to specific subcellular membrane compartment(s) to sustain their spatially restricted distribution, whether dynamic or static- to recognize membrane-associated substrates, is unclear.

One mode of protein targeting to membrane compartments is through post-translational lipid modifications^3,4^. Prenylation, a type of lipid modifications, is essential for the prenyl-protein association with the endo and peripheral membrane compartments, as exemplified by small GTPase family members^5,6^. Four known mammalian prenyltransferases (PTs), Ftase and GGtase 1-3, enable protein prenylation of a large class of proteins that contain a cysteine residue near the C-terminus, usually part of the canonical CaaX motif signature (C: cysteine, a: aliphatic, X: any amino acid) and atypical CaaX motifs (CxxX, CCxX, xCCX, xxCC, xCxC and CCxxX) ^7^. PTs during a prenylation reaction catalyze the attachment of either a farnesyl (15-carbon) or geranylgeranyl (20-carbon) isoprenoid lipid *via* irreversible thioether linkage to the cysteine residue in the CaaX motif of the substrate prenyl-protein^8,9^. Moreover, prenylated-CaaX motifs may undergo further processing, which is believed to increase hydrophobicity, thereby enhancing the membrane association of prenylated proteins ^6,10^. Although, prenylation is essential for target protein-membrane association, it alone does not ensure the correct delivery of CaaX-proteins to specific subcellular membrane compartments ^6,10^. For example, CaaX-proteins that are delivered to the PM generally contain what are termed as second signal sequences, usually juxtaposed to CaaX motifs, which undergo reversible palmitoylation (HRAS, NRAS, RHOB, TC10, and FBXL2) or polybasic amino acid residues (KRAS)^11,12^. The subcellular distribution of prenyl-proteins is also governed by their trafficking factors. For example, cytosolic factor δ subunit of type 6 phosphodiesterase (PDE6δ) possesses the structural capability to accommodate isoprenoid lipid groups into its hydrophobic pocket shielding the prenyl-lipid in the aqueous milieu of cytoplasm allowing the transport of the prenyl-protein from one membrane to another membrane compartment ^13,14^.

Mitochondria are distinctive double membranous organelles with their own genome (mtDNA), and as such, mitochondria-driven mechanisms impact a wide spectrum of cellular processes including cell bioenergetics, metabolism, viability, proliferation, and differentiation^15^. Remarkably, mitochondria are highly plastic organelles organized in dynamic networks that undergo constant morphological remodeling (fusion/fission cycles, intra-cell mobility) in response to cues and stressors for efficient homeostatic cellular function. Abnormal mitochondrial dynamics compromise the functional efficiency of re-modeled mitochondrial networks, triggering mitophagy wherein damaged, depolarized, fragmented, and inefficient mitochondria are selectively engulfed by autophagosomes and trafficked to lysosomes for degradation ^16^. However, such destruction and recycling of whole mitochondria is not suitable for the targeted regulation of specific mitochondrial proteins. While basal mitophagy ensures an efficient pool of healthy mitochondria, persistent mitophagy may be counterproductive. The mitochondrial proteome (∼1100 proteins) is encoded primarily by the nucleus (>98%) and distributed between the outer mitochondrial membrane (OMM), the inner mitochondrial membrane (IMM), and the two aqueous spaces created between the two membranes, intermembrane space (IMS) and mitochondrial matrix (MM). Nuclear DNA encoded mitochondrial proteins are targeted to each of these compartments primarily by mitochondria targeting sequences (MTS) integral to polypeptides and mitochondria specific chaperones, such as HSP90, that dock on OMM receptors to deliver the clients for distribution within various sub-compartments^17,18^. Although, for wholesale clearance of damaged mitochondria, E3s such as Parkin and FBXL4 are important to promote and/or suppress mitophagy by non-selective ubiquitylation of OMM proteins and specific mitophagy/autophagy receptors^19,20^, however, the UPS-regulated selective protein ubiquitylation at each mitochondrial sub-compartment for fine-tuned mitochondria-driven homeostatic functions without triggering mitophagy is unclear. Here we report FBXO10 is a geranylgeranylated-CaaX-protein-E3, functional at outer mitochondrial membrane (OMM). We provide mechanistic insights into why, despite lacking a canonical MTS, FBXO10 is specifically distributed at OMM, but not to other membrane compartments. Leveraging redistribution of geranylgeranylation-deficient FBXO10(C953S) away from the OMM, we exposed potential FBXO10 substrates at OMM by comparative quantitative mass spectrometry analyses of enriched mitochondria. We show that phosphoglycerate mutase 5 (PGAM5) is ubiquitylated and targeted for degradation *via* FBXO10, but not FBXO10(C953S), at OMM, during the myogenic differentiation program. Supplemented with FBXO10 loss-of-function studies, we delineate mechanistic basis for the dysregulation of mitochondrial homeostasis and mitochondria-driven impaired myogenic differentiation by geranylgeranylation-deficient FBXO10(C953S) in human iPSCs and murine myoblasts. Thus, our studies demonstrate the critical importance of the maintenance of dynamic OMM distribution of FBXO10 for the selective OMM proteostasis and function.

## Results

### FBXO10 maintains mitochondrial proteostasis

F-box proteins (69 in humans) are the substrate receptors of CRL1 (CUL1-Ring protein Ligase) complexes (aka SCF for SKP1, CUL1, F-box protein complexes)^1,2^. A close inspection of the coding sequences of FBXO10 and its paralog FBXO11 revealed that FBXO10, but not FBXO11 which is a nuclear protein ^21^, terminates with an atypical CaaX motif sequence highly conserved across species orthologs with a cysteine residue at position 953 (C953) (Fig. 1A and Fig. S1A). CaaX motif containing proteins, upon prenylation, are known to associate with cellular membrane compartments^8,9^. Therefore, we hypothesized that FBXO10 is a membrane-associated F-box protein and sought to investigate the CaaX motif dependent subcellular distribution of FBXO10 with various orthogonal assays utilizing live cell confocal microscopy, subcellular fractionation, and fluorescence-activated cell sorting (FACS). First, we swapped the C953 residue to serine (C953S) with site-directed mutagenesis to generate mutant FBX010 that lacks an intact CaaX motif [FBXO10(C953S)]. This serves as a genetic tool to interrogate the CaaX motif dependent subcellular distribution and functions of FBXO10 (Fig. S1B). The live-cell confocal imaging with organelle specific markers and colocalization quantification analysis demonstrated that GFP-FBXO10 decorates the mitochondrial network (Fig. 1B panel 1, Fig. 1K top panel and Fig. S1C panel 1). In contrast, mutation of cysteine 953 in the CaaX motif rendered GFP-FBXO10(C953S) cytosolic as revealed by a homogeneous fluorescence with negatively visualized organelles (Fig. 1B panel 2, Fig. 1K bottom panel and Fig. S1C panels 2-4). To monitor the plasma membrane (PM) and motile vesicular compartment (VC), we designed a new fluorescent-probe PMV-SeQ (Fig. S1D) based on juxtaposed palmitoylation and prenylation signal motifs identified by our recent work with another PM-localized E3-SCF^FBXL2^ ^11^. Investigation with the PMV-SeQ confocal microscopy demonstrated that GFP-FBXO10 and GFP-FBXO10 (C953S) do not decorate PM and VC (Fig. 1B, panel 3 and Fig. S1C panel 3). Analysis of endoplasmic reticulum (ER) compartment revealed insignificant ER specific localization indistinguishable between GFP-FBXO10 and CaaX mutant GFP-FBXO10(C953S) (Fig. 1B panel 4 and Fig. S1C panel 4). Subcellular fractionation of cells expressing Strep-FBXO10 and Strep-FBXO10(C953S) validated the exclusive enrichment of FBXO10 in mitochondrial fractions (Fig. 1C, lane 2 vs 5, top panel). In contrast, Strep-FBXO10(C953S) enriched in soluble cytosolic fractions but not in mitochondria (Fig. 1C, lane 3 vs 6, top panel and Fig. S1J), consistent with its homogenous cellular distribution pattern (Fig. 1B). Flow cytometry-based quantification of FBXO10 fluorescence in intact mitochondrial fractions isolated from cells expressing ptd-Tomato-FBXO10 and ptd-Tomato-FBXO10(C953S) further confirmed that whereas FBXO10 is associated with mitochondria, the CaaX-mutant FBXO10(C953S) is delocalized away from mitochondria (Fig. 1D). Next, we investigated if FBXO10 is able to assemble into a multi-subunit CRL complex at mitochondria. Immunoprecipitation of FBXO10 from enriched mitochondrial fractions demonstrated the co-immunoprecipitation of SKP1 and CUL1 (Fig. S1E) suggesting FBXO10 forms an authentic CRL1 associated with mitochondria. Since the CaaX-mutant FBXO10(C953S) is delocalized away from mitochondria (Fig. 1B-D and Fig. S1C), we carried out immunoprecipitations from whole cell lysates and found that SKP1 and CUL1 also co-immunoprecipitated with FBXO10(C953S). However, we noticed a detectable difference in neddylation of co-immunoprecipitated CUL1 when compared to wild-type FBXO10 (Fig. 1E). This result is consistent with the notion that FBXO10 may engage substrates, which promote cullin neddylation^22,23^, for ubiquitylation and subsequent proteasomal degradation at mitochondria.

**Figure 1.**
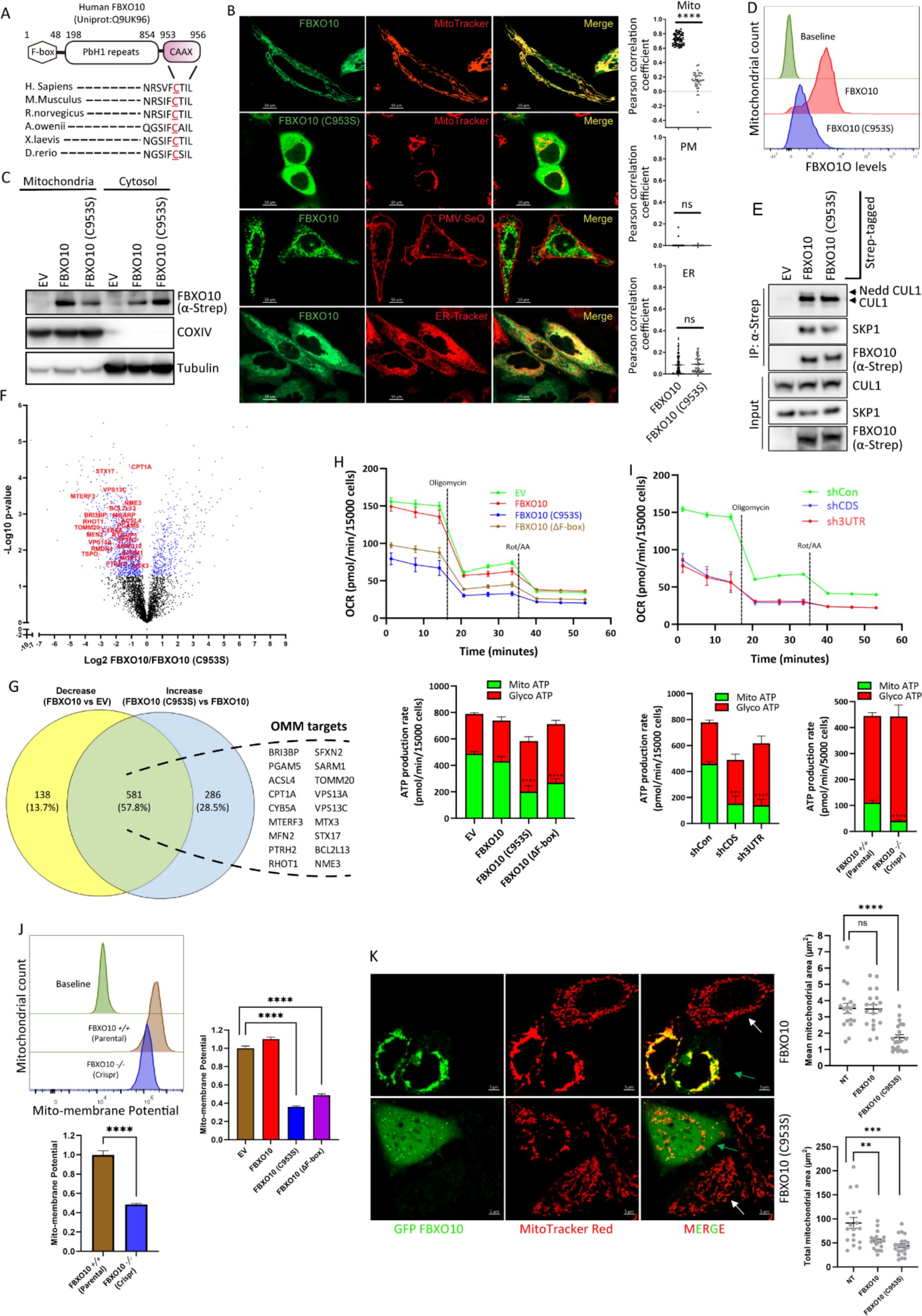
Mitochondrial function and selective proteostasis at outer mitochondrial membrane (OMM) controlled by FBXO10. **(A)** *FBXO10 polypeptide terminates in a CaaX motif with Cysteine953 conserved across species.* FBXO10 schematic at top shows the predicted protein domain configuration highlighting F-box, PBH1 and newly identified C-terminal CaaX motif. The alignment of C-terminal amino acid sequences of FBXO10 orthologs is shown. CaaX motif with Cysteine953 is underlined. The numerals on top represent the position from the N-terminus corresponding to human polypeptide. **(B)** *FBXO10 subcellular distribution at mitochondria depends on the integrity of CaaX-Cysteine953.* HeLa cells expressing GFP-FBXO10 (panels 1, 3 and 4) and CaaX-motif mutant GFP-FBXO10(C953S) (panel 2) were either treated with MitoTracker, ER-Tracker or co-transfected with PMV-SeQ to decorate mitochondrial networks (panels 1 and 2), ER (panel 4) and plasma membranes (PM, panel 3), respectively. ER-Tracker and Mito-Tracker were added 15 minutes prior to live cell confocal imaging by ZEISS-LSM700 confocal microscope, as shown by representative captured images. Pearson correlation coefficients (PCC) for the subcellular organelle colocalization of GFP-FBXO10 and GFP-FBXO10 (C953S) were analyzed in different cells [MitoTracker: N= 64 for FBXO10 and N= 35 for FBXO10 (C953S); PMV-SeQ: N= 85 for FBXO10 and N= 70 for FBXO10 (C953S); ER-Tracker : N= 157 for FBXO10 and N= 33 for FBXO10 (C953S)]. Statistical analysis of plotted PPCs in scatter plots confirmed subcellular distribution of GFP-FBXO10 exclusively at mitochondria in contrast to CaaX-motif mutant GFP-FBXO10(C953S) which re-distributes away from the mitochondria, ER and PM. p-values were calculated by student-T-test. Scale bar 10 µM. **(C)** *FBXO10, but not CaaX-mutant FBXO10(C953S), is retained in subcellular mitochondrial fractions.* STREP-tagged FBXO10, FBXO10(C953S), and vehicle (EV) were expressed in HEK-293T cells and post-harvesting cell lysates were subjected to subcellular fractionation using stepwise ultracentrifugation (see Methods). Enriched mitochondrial and cytosolic fractions were assayed by SDS-PAGE and immunoblotted as indicated. Representative of three independent experiments is shown. **(D)** *CaaX-mutant FBXO10(C953S) is delocalized away from mitochondria.* Enriched mitochondrial fractions were isolated from HEK293T cells expressing ptd-Tomato-FBXO1O, ptd-Tomato-FBXO10 (C953S) and non-expressing controls by stepwise centrifugation. Isolated mitochondrial fractions were incubated with MitoView green to allow mitochondrial tracking during flowcytometry analysis (FACS) for the quantification of ptd-Tomato-FBXO10 and ptd-Tomato-FBXO10 (C953S) signals. Representative FACS histogram confirms localization of FBXO10 at mitochondria (middle panel) shown as rightward shifted peak from the baseline non-expressing controls (top panel) and delocalization of FBXO10 (C953S) away from mitochondria shown as peak closer/matching baseline non-expressing controls (bottom panel). Representative of three independent experiments is shown. **(E)** *FBXO10 and CaaX-mutant FBXO10(C953S) assemble into Cullin1-Ring-Ligase (CRL-1) complexes.* STREP-tagged FBXO10, FBXO10(C953S) and empty vector (EV) were expressed in HEK293T cells and post cell harvesting, STREP-Tactin immunoprecipitations were performed with standard protocols. Subsequently, immunoprecipitated complexes were subject to SDS-PAGE and immunoblotting as shown. Shown is representative of three independent experiments. **(F)** *Comparative mass spectrometry uncovers selective mitochondrial proteostasis via FBXO10.* Perturbations in mitochondria-associated proteome was assayed by label free quantification (LFQ) mass spectrometry analysis (LFQ-MS/MS) of enriched mitochondrial fractions isolated by stepwise centrifugation from stable HEK293T cells expressing FBXO10, FBXO10 (C953S) and empty vector. Computational analysis of detected data sets under the three experimental conditions revealed changes in mitochondria-associated protein levels. The volcano plot shows significantly altered proteins between FBXO10 and FBXO10(C953S) data sets. OMM proteins that showed significant reciprocal protein level changes, i.e., decrease upon FBXO10 but increase upon FBXO10(C953S) expression, are highlighted (≥2 folds, at FDR 5%). FDR: False discovery rate. Three independent biological replicates of each FBXO10, FBXO10 (C953S) and empty vector samples were assayed by LFQ-MS/MS. **(G)** *Candidate OMM targets of FBXO10 revealed by analysis of* LFQ-MS/MS *data sets.* Computational analysis of data detected by LFQ-MS/MS under the experimental conditions mitochondrial proteome revealed changes in mitochondria-associated protein levels. Venn diagram depicts overlap of significantly deregulated protein numbers, i.e., decreased in FBXO10 *vs* EV and increased in FBXO10 (C953S) *vs* FBXO10 data sets. List of eighteen OMM proteins with reciprocal protein level changes, i.e., decrease upon FBXO10 but increase upon FBXO10(C953S) expression is shown (≥2 folds, at FDR 5%). Three independent biological replicates of each FBXO10, FBXO10 (C953S) and empty vector samples were assayed by LFQ-MS/MS. **(H)** *CaaX-mutant FBXO10(C953S) and E3-ligase activity deEicient FBXO10(ΔF-box) inhibit mitochondrial ATP production.* Measurement of mitochondrial ATP production was carried out in C2C12 cell stably expressing FBXO10, FBXO10 (C953S), FBXO10 (Δ-Fbox) and empty vector (EV) by Seahorse XF Real-Time ATP Assay. Line graphs show oxygen consumption rate (OCR) and bar graph shows mitochondrial and glycolytic ATP production rate (pmol/min) obtained from OCR and ECAR measurements. Bar graphs represent quantifications of ten biological replicates. Error bars indicate SEM. **(I)** *FBXO10 gene deletion and silencing inhibit mitochondrial ATP production.* CRISPR/Cas9-mediated FBXO10 gene deletion was carried out in C2C12 cells. FBXO10 silencing was carried out with two independent shRNAs targeted to untranslated (3’UTR) and coding (CDS) regions in C2C12 cells. Measurement of mitochondrial ATP production rate in indicated samples (controls, shRNA and CRISPR-Cas9 clone), was performed by Seahorse XF Real-Time ATP Assay as in (H). n= 10 for shCon, sh3UTR and shCDS; n= 11 for FBXO10 *+/+*, n=9 for FBX010 *-/-* where n represents the number of biological replicates. **(J)** *FBXO10 gene deletion, CaaX-mutant FBXO10(C953S) and E3-ligase activity deEicient FBXO10(ΔF-box) inhibit mitochondrial membrane potential.* Measurement of mitochondrial membrane potential was carried out by flowcytometry upon TMRM treatment (100nM) in differentiated C2C12 cell stably expressing FBXO10, FBXO10(C953S), FBXO10(Δ-Fbox), empty vector (EV) and CRISPR/Cas9-mediated FBXO10 gene deleted C2C12 clone (bottom). Bar graphs represent quantifications of three biological replicates for EV, FBXO10, FBXO10(C953S), FBXO10(Δ-Fbox) four biological replicates for FBXO10 *+/+*, n=9 for FBX010 *-/-.* p-values were calculated by a student-T-test. Error bars indicate SEM. **(K)** *FBXO10 promotes perinuclear mitochondrial clustering whereas CaaX-mutant FBXO10(C953S) promotes mitochondrial fragmentation.* HeLa cells expressing GFP-FBXO10 (top panel) and GFP-FBXO10(C953S) (bottom panel) were treated with MitoTracker for ∼30 minutes prior to live cell confocal imaging using ZEISS-LSM750 microscope. Representative images depict the changes in mitochondrial network dynamics upon FBXO10 and FBXO10 (C953S) expression. Green arrows point to transfected cells whereas white arrows indicate non-transfected cell in the same field of view analyzed for morphological dynamic alterations. Scatter plots show total mitochondrial network areas/cell and mean mitochondrial areas calculated from non-transfected (NT), FBXO10 and FBXO10 (C953S) expressing cells using ImageJ mitochondrial analyzer software. [n=18 NT, n=19 FBXO10, n=21 FBXO10 (C953S)], p-values were calculated by a student-T-test. Error bars indicate SEM. Scale bar, 5 µm.

Because CaaX mutant FBXO10(C953S) delocalizes away from mitochondria (Fig. 1B-D and Fig. S1C) whereas FBXO10 enriches as an authentic CRL1 at mitochondria (Fig. 1B-D, Fig. S1C and Fig. 1E), we exploited the delocalized FBXO10(C953S) mutant to investigate the importance for the maintenance of the subcellular distribution on the substrate degradation at a specific subcellular membrane organelle, mitochondria. Accordingly, we analyzed perturbations in the mitochondria-associated proteome by carrying out label-free quantitative mass spectrometry analysis (LFQ-MS/MS) of enriched mitochondrial protein fractions purified from doxycycline-inducible cells stably expressing FLAG-FBXO10, FLAG-FBXO10(C953S), and empty vector (EV), at low or near endogenous levels controlled *via* doxycycline dosage (Figs. S1F and S1G). Proteins whose abundance was reciprocally impacted, i.e., increased by FBXO10 (C953S) expression but decreased upon FBXO10 expression, were considered as potential targets of FBXO10, either direct or indirect. Our unbiased quantitative mass spectrometry data unmasked the role of FBXO10 in selective proteostasis at mitochondria and revealed dysregulated proteostasis of mitochondria-associated proteins by the expression of CaaX-mutant FBXO10(C953S) (Fig. 1F, 1G, S1H and S1I; LFQ data available on request). As we found FBXO10 resides in outer mitochondrial membrane (OMM) but not IMM or MM (refer: Fig. 2F, G), we reasoned that direct substrates of FBXO10 should also localize to the OMM. Therefore, we filtered and ranked the identified proteins based on their OMM and/or OMM-associated localization with a hypothesis that FBXO10 more likely has direct access to OMM and/or OMM-associated protein targets for ubiquitylation to promote subsequent proteasomal degradation. We discovered eighteen OMM and/or OMM-associated protein targets that showed significant reciprocal changes in protein levels (≥2-fold change, 5% FDR) (Fig. 1F and G; LFQ data available on request). We validated ubiquitylation and degradation of phosphoglycerate mutase 5 (PGAM5) *via* FBXO10 (refer: Fig. 6 and Fig. S6). Moreover, functional annotation clustering analysis of the significantly dysregulated proteins (FDR 5%, deregulation ≥ 2 folds identified by MS analysis) indeed showed the highest enrichment score for mitochondria, as expected, and revealed that FBXO10-mediated protein turnover at the OMM impacts a range of cellular processes such as protein translation, carbohydrate metabolism, oxidative phosphorylation, and mRNA processing. (Fig. S1I).

**Figure 2.**
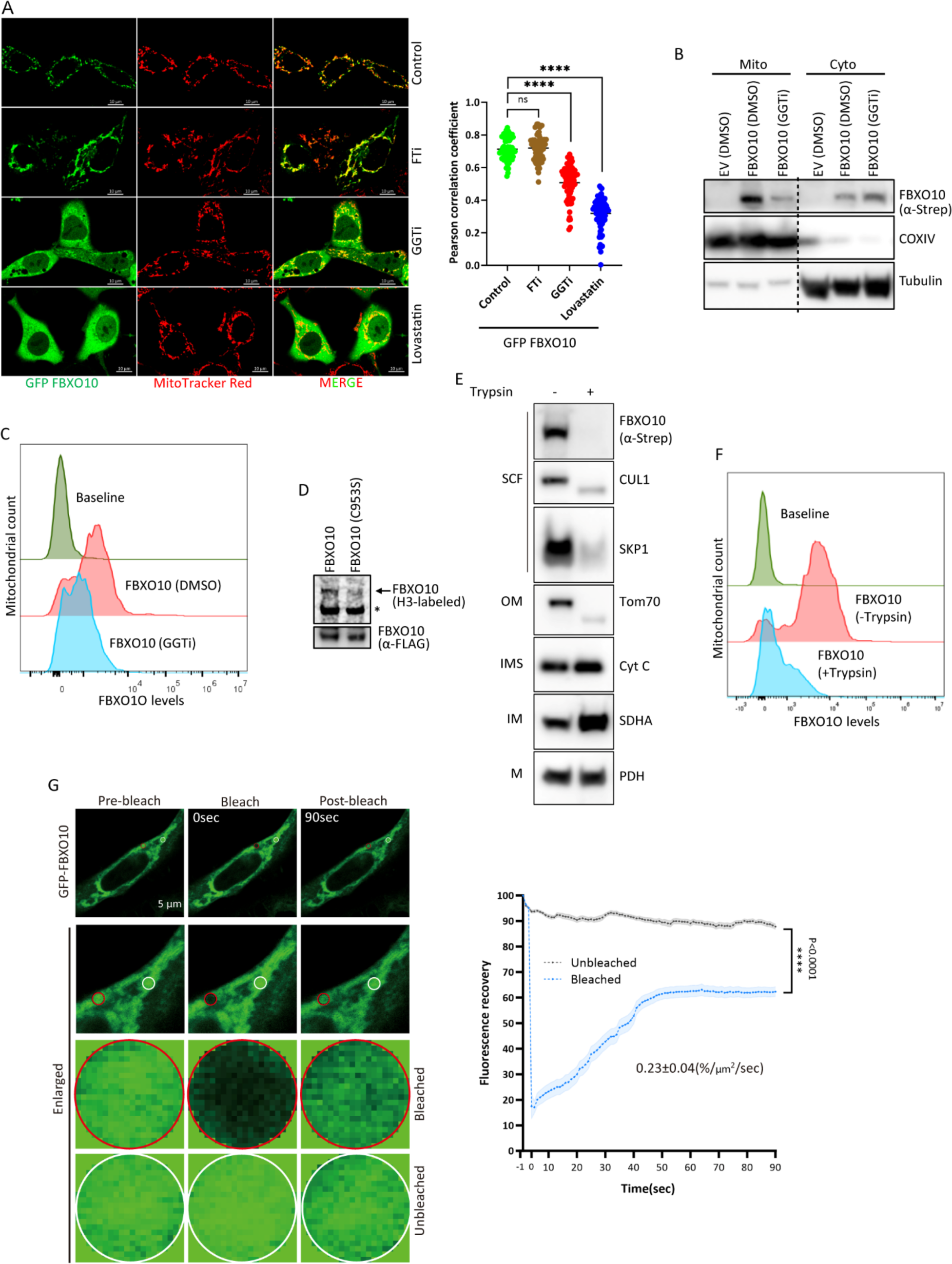
Geranylgeranylation of CaaX-cysteine953 is indispensable for the distribution of FBXO10 at outer mitochondrial membrane. **(A)** *Inhibition of geranylgeranylation re-distributes GFP-FBXO1O away from mitochondria into cytosol.* HeLa cells expressing GFP-FBXO10 were treated with geranylgeranylation inhibitor (GGTi-2418, 50 µM) farnesylation inhibitor (FTi-lonafarnib, 10 µM), clinical drug statin (lovastatin, 15 µM) and vehicle for 16 hours. MitoTracker was added to decorate mitochondrial for ∼30 minutes prior to live cell confocal imaging using ZEISS-LSM700 microscope. Representative images depict the changes in mitochondrial distribution of GFP-FBXO10 (panel 1) upon inhibition of farnesylation (panel 2), geranylgeranylation (panel 3) and mevalonate pathway (panel 4). Pearson correlation coefficients (PCC) for the colocalization of GFP-FBXO10 and mitochondria were analyzed in different cells [vehicle (n=70), GGTi (n=72), Fti (n=73) and lovastatin (n= 77)]. Statistical analysis of plotted PPCs in scatter plots is shown (right). p-values were calculated by student-T-test. Scale bar 10µM. **(B)** *Inhibition of geranylgeranylation results in loss of FBXO10 from subcellular mitochondrial fractions.* HEK-293T cells expressing STREP-tagged FBXO10 cells were treated with geranylgeranylation inhibitor (GGTi-2418, 50 µM) or vehicle DMSO for 16 hours. Post-harvesting cell lysates were subjected to subcellular fractionation using stepwise ultracentrifugation (see Methods). Enriched mitochondrial and cytosolic fractions were assayed by SDS-PAGE and immunoblotted as indicated. Shown is representative of two independent experiments. **(C)** *GGTi-2418 delocalizes FBXO10 from mitochondria.* HeLa cells expressing ptd-Tomato-FBXO10 were treated with geranylgeranylation inhibitor (GGTi-2418, 50 µM) or vehicle DMSO for 16 hours. Enriched mitochondrial fractions were isolated by stepwise centrifugation. Isolated mitochondrial fractions were incubated with MitoView green facilitating mitochondrial tracking during flowcytometry analysis (FACS) for the quantification of mitochondria-associated ptd-Tomato-FBXO10 signal. Representative FACS histogram confirms localization of FBXO10 at mitochondria (middle panel) shown as rightward shifted peak from the baseline non-expressing controls (top panel) and delocalization of FBXO10 away from mitochondria upon GGTi-2418 treatment shown as peak closer/matching baseline non-expressing controls (bottom panel). Shown is representative of three independent experiments. **(D)** *Metabolic labelling with tritiated mevalonate shows Cystine953 acceptor site for the FBXO10 prenylation.* HEK293T cells expressing FLAG-FBXO10 and FLAG-FBXO10(C953S) were metabolically labeled with tritiated (H^3^)-mevalonolactone according to the protocol described in the methods section. Post-labeling, cells were harvested, and whole cell lysates were processed to carry out anti-FLAG immunoprecipitations. Tritium (H^3^) labeled immunocomplexes and lysates were subjected to SDS-PAGE using pre-cast gradient Bolt^TM^ 4-12% Bis-Tris Plus gels and transferred to a PVDF membrane. Subsequently, the PVDF membrane was exposed to a Phosphor-screen and H^3^-labeled proteins were detected with the Phosphor-imager. The autoradiograph (top), developed after exposure to Phosphor-screen, shows that FLAG-FBXO10, but not FLAG-FBXO10(C953S) is prenylated. (*) indicates non-specific. Bottom: PVDF membrane was probed with anti-FLAG immunoblotting. **(E)** *FBXO10 along with SCF-subunits associate with outer mitochondrial membrane (OMM).* Enriched intact mitochondrial fractions were isolated from HEK293T cells expressing STREP-FBXO1O by stepwise centrifugation. Isolated mitochondrial fractions were subjected to Trypsin protease protection (TPP) assay protocol (see methods). Mitochondrial lysates prepared from samples that underwent TPP-assay and untreated controls were subjected to SDS-PAGE followed by immunoblotting for the SCF-FBXO10 components and sub-mitochondrial membrane compartment specific markers, as indicated. Representative of three independent experiments is shown. **(F)** *FBXO10 localizes at outer mitochondrial membrane (OMM).* Enriched intact mitochondrial fractions were isolated from HEK293T cells expressing GFP-FBXO1O by stepwise centrifugation. Isolated mitochondrial fractions were subjected to Trypsin protease protection (TPP) as in (H). Samples that underwent TPP-assay and untreated controls were subjected to flowcytometry analysis (FACS) to quantify mitochondria-associated GFP-FBXO10 signal. Representative FACS histogram confirms FBXO10 at OMM (middle panel) shown as rightward shifted peak in untreated samples from the baseline non-expressing controls (top panel) and its loss from OMM due to Trypsin protease access at OMM shaving off GFP-FBXO10 in TPP-samples shown as leftward shifted peak closer/matching baseline non-expressing controls (bottom panel). Representative of three independent experiments is shown. **(G)** *Dynamic subcellular distribution of FBXO10 at mitochondria.* Fluorescence recovery after photobleaching (FRAP) analysis was carried out to monitor mobility of FBXO10 in HeLa cells expressing GFP-FBXO10. After focal photodestruction of the GFP signal with 488nm laser, the recovery of the GFP-FBXO10 signal in the bleached area was recorded over indicated time period using ZEISS-LSM700 microscope. In parallel an unbleached focal area was also monitored as control. Shown are the representative images of 18 independent cells analyzed for FRAP. The plot shows averaged GFP-FBXO10 signal recovery of 18 independent bleached areas from separate cells. GFP-signal before photodestruction was set as 100%. Calculated Recovery Rate: 0.23±0.04(%/µm2/sec). Error bar: SEM

### CaaX-mutant FBXO10(C953S) and FBXO10 loss dysregulate mitochondrial ATP production, membrane potential, and morphological dynamics

To examine the functional consequences of impaired FBXO10 subcellular distribution and dysregulated protein turnover at the OMM, we investigated mitochondrial ATP production, mitochondrial membrane potential, and mitochondrial morphological dynamics. In addition to FBXO10(C953S), we also generated FBXO10(ΔF-box) that is unable to assemble into a functional SCF yet able to localize correctly at mitochondria (Fig. S1J and S1K) and hence expected to sequester substrates away from proteasomal degradation ^24^. The rate of mitochondrial and glycolytic ATP production was measured by Seahorse XF real-time ATP rate assay in cells stably expressing wild type-FBXO10, delocalized-FBXO10(C953S), and mitochondria-localized but E3-ligase activity deficient-FBXO10(ΔF-box). We found that mitochondrial ATP production was inhibited by delocalized-FBXO10(C953S) and FBXO10(ΔF-box) expression, but not by wild type-FBXO10 (Fig. 1H), suggesting that FBXO10(C953S) and FBX019(ΔF-box) act in a dominant negative manner. These results suggest that the dysregulation of OMM proteolysis either by delocalizing FBXO10 away from mitochondria and/or by an incompetent-ligase FBXO10(ΔF-box) at mitochondria impairs mitochondrial ATP production. Interestingly, we found proportionally glycolytic ATP content increased in cells expressing delocalized-FBXO10(C953S) and FBXO10(ΔF-box) compared to cells expressing FBXO10 and empty vector control (Fig. 1H). Next, we examined the loss-of-function phenotype of FBXO10, using CRISPR-Cas9 gene editing and/or depletion with two different short hairpin RNAs (shRNAs) targeted at coding sequence (CDS) and 3’ untranslated region (3’ UTR) (Fig. S6E and Fig. S1M). Our results demonstrated that both FBXO10 deletion and/or depletion inhibited mitochondrial ATP production but enhanced the proportion of glycolytic ATP (Fig. 1I), consistent with the results *via* the expression of delocalized-FBXO10(C953S) and FBXO10(ΔF-box) (Fig. 1H). To measure mitochondrial membrane potential, we carried out TMRM staining (a fluorescent probe to quantify mitochondrial membrane potential) in cells stably expressing wild type-FBXO10, delocalized-FBXO10(C953S),) mitochondria-localized but E3-ligase activity deficient-FBXO10(ΔF-box), and a CRISPR-Cas9 edited FBXO10 gene deleted clone. We found mitochondrial membrane potential (MPP) was inhibited by delocalized-FBXO10(C953S) and FBXO10(ΔF-box) expression, but not by wild type-FBXO10 (Fig. 1J, and Fig. S1L). FBXO10 gene deletion also inhibited MPP (Fig. 1J), consistent with expression of delocalized-FBXO10(C953S) and/or FBXO10(ΔF-box). These results show that the dysregulation of OMM proteolysis results in loss of mitochondrial membrane potential indicating mitochondrial damage upon expression of delocalized-FBXO10(C953S) and FBXO10(ΔF-box). Taken together, our results show that the loss of FBXO10 (deletion or silencing), delocalization of FBXO10 away from mitochondria, and incompetent-FBXO10 ligase at mitochondria inhibits mitochondrial ATP production and mitochondrial membrane potential.

Next, we investigated the impact on the mitochondrial morphological dynamics (i.e., fusion/fission) with live cell confocal microscopy upon re-distribution of FBXO10 away from mitochondria and FBXO10 silencing. To this end, we analyzed mitochondrial network morphological changes in cells expressing delocalized-FBXO10(C953S) compared to wild type-FBXO10 expressing and non-transfected (NT) cells. Wild type FBXO10 promoted hyperfused spaghetti-like mitochondrial networks and clustering of mitochondria around perinuclear regions compared to non-transfected cells under same conditions (Fig. 1K top panel, Fig. 1B panel 1 and Fig. S1C panel 1). Quantification revealed significant reduction of total mitochondrial area occupied per cell when FBXO10 was expressed, consistent with clustered and/or hyperfused mitochondrial mass (Fig. 1K, left scatter plot). In contrast, mitochondrial networks in FBXO10(C953S)-expressing cells were visualized as largely non-reticulate, scattered, and irregular networks, apparently revealing mitochondrial fragmentation (Fig. 1B panel 2, Fig. 1K, bottom panel and Fig. S1C panel 2) compared to typical reticulate, well-spread tubular networks in non-transfected cells (Fig. 1K, refer to white arrows). Quantitative measurements and analysis show significant reduction in mean mitochondrial area upon expression of delocalized-FBXO10(C953S) compared to non-transfected cells as well as wild-type FBXO10-expressing cells (Fig. 1K, right scatter plot) reflective of a fragmentation phenotype. Interestingly, measurements of mitochondrial membrane potential (MMP) show loss of MMP upon expression of delocalized-FBXO10(C953S) consistent with fragmented and damaged mitochondria (Fig. 1J). Furthermore, we visualized intact mitochondrial networks by a fluorescent probe retained within the intact mitochondria (MitoTracker Red) upon FBXO10 silencing with two different shRNAs targeting CDS and 3’ UTR. While in non-targeting shRNA treated cells mitochondria appeared as typical reticulate networks, FBXO10 silencing showed reduction in reticulate mitochondria content appearing as dispersed consistent with mitochondrial damage and fragmentation (Fig. S1M). Loss of mitochondrial potential and fragmentation of mitochondria upon expression of delocalized-FBXO10(C953S) and/or FBXO10 depletion is consistent with the reduction of mitochondrial ATP production seen under these conditions. Thus, our results show functional FBXO10 subcellar mitochondrial distribution, dependent on its intact cysteine953 in C-terminal CaaX motif (Fig. 1B-D and Fig. S1C), is critical for the homeostatic mitochondrial proteostasis (Fig. 1F-G and Fig. S1H-I) and mitochondrial function (Fig. 1H-K).

### FBXO10 is geranylgeranylated on Cysteine953 for dynamic subcellular distribution to outer mitochondrial membrane (OMM)

The majority of CaaX-motif containing proteins undergo post-translation CaaX modification, prototypically described in RAS oncoproteins of small GTPase family, with prenyl isoprenoids (i.e., either 15-carbon farnesyl or 20-carbon geranylgeranyl) followed by prenyl-CaaX motif processing at Golgi/ER endomembranes, enabling dynamic association with membrane compartments^8,9^. Human FBXO10 contains an atypical CaaX motif, *CTIL*, conserved across species orthologs with cysteine953 (Fig. 1A), essential for its localization to mitochondria (Fig. 1A-C). Therefore, we tested the hypothesis that cysteine953 is a prenyl isoprenoid acceptor site in FBXO10 and sought to determine whether FBXO10 is a candidate for farnesylation and/or geranylgeranylation. To this end, we first treated GFP-FBXO10 expressing cells with farnesylation and geranylgeranylation inhibitors (FTi and GGTi) and visualized the subcellular mitochondrial distribution with live cell confocal microscopy. We found treatment with GGTi-2418, but not FTi-lonafarnib, re-distributed GFP-FBXO1O away from mitochondria into cytosol (Fig. 2A), phenocopying the cellular distribution shown by cysteine953 mutant GFP-FBXO1O(C953S) (Fig. 1B), as homogeneous fluorescence with negatively visualized organelles (Fig. 1B and Fig. 2A panel 3). Interestingly, we found that the treatment with lovastatin (statins are widely used drugs in clinics to lower blood cholesterol by inhibiting a key step in cellular mevalonate pathway)^25^ leads to the redistribution of GFP-FBXO1O away from mitochondria into cytosol, suggesting FBXO10 is a downstream target of statins (Fig. 2A panel 4). Inhibition of the cellular mevalonate pathway by statins prevents the production of geranylgeranyl isoprenoids. Subcellular fractionation of FBXO1O to enriched mitochondria was inhibited by GGTi-2418 treatment, akin to the delocalized cysteine953 mutant FBXO1O(C953S) (Fig. 1C and 2B). Moreover, quantification of FBXO10 by flow cytometry (FACS) in intact mitochondrial fractions further confirmed that GGTi-2418 treatment, but not FTi-lonafarnib, delocalized FBXO10 away from the mitochondria (Fig. 2C). These results demonstrate that FBXO10 undergoes geranylgeranylation, but not farnesylation, for the subcellular mitochondrial distribution, and suggested that cysteine953 is an acceptor site for the geranylgeranylation. To test directly if FBXO10 is indeed geranylgeranylated dependent on CaaX motif cysteine953, we carried out metabolic labeling of mammalian cells with tritiated [H^3^]-mevalonate, and directly analyzed incorporation of label into immunoprecipitated FBXO10 and cysteine953 mutant FBXO1O(C953S) by autoradiography. Our results show that whereas FBXO10 is geranylgeranylated, the cysteine953 mutant FBXO1O(C953S) completely lacked detectable geranylgeranylation signal (Fig. 2D).

Structurally, the mitochondrion is a double membranous organelle composed of OMM, IMM, and the two aqueous spaces, IMS and matrix. Each of these compartments perform specialized tasks due to distinctive protein signatures. To determine the exact sub-mitochondrial localization of FBXO10, we performed a trypsin protease protection assay in which we exposed intact double membranous mitochondria expressing Strep-FBXO10 to trypsin protease added exogenously. In this assay, the intact mitochondrion is expected to protect IMM, IMS, and matrix proteins from trypsin digestion, whereas proteins associated with OMM are accessible to trypsin protease for digestion. We found, just like OMM marker protein TOM70, the Strep-FBXO10 along with its SCF cognate subunits SKP1 and Cullin1 were shaved off from the intact mitochondria by trypsin digestion, whereas protein markers of IMM (SDH), IMS (cytochrome C), and matrix (PDH) remained refractory to trypsin protease digestion (Fig. 2E). These results demonstrate that geranylgeranylated-FBXO10 resides at the OMM. Furthermore, we confirmed the OMM targeting of geranylgeranylated-FBXO10 by quantifying the extent of trypsin protease shaving in isolated mitochondria-expressing flourescent-ptd-Tomato-FBXO10 by flow cytometry and found that the entire peak of flourescent-FBXO10 was sensitive to trypsin digestion rendering it to baseline (Fig. 2F). To examine if geranylgeranylated-FBXO10 associates with OMM in a permanent or dynamic distribution pattern, we carried out fluorescence recovery after photobleaching (FRAP) of GFP-FBXO10 at mitochondria (Fig. 2G and Fig. S2A). After photodestroying fluorescent GFP-FBXO10 focally at mitochondria, we found that the GFP-FBXO10 fluorescence recovered at the bleached mitochondrial spots with a calculated recovery rate of 15.78±2 (%/µm2/msec), revealing a dynamic trafficking pattern of geranylgeranylated-FBXO10. Moreover, the quantification of the FRAP data revealed that the recovered fluorescence of GFP-FBXO10 plateaued at significantly lower levels after photobleaching (Fig. 2G), which suggests that fluid phase diffusion may not be the sole means to account for the FBXO10 trafficking to mitochondria. Taken together, our results demonstrate that cysteine953 is the geranylgeranyl acceptor for the dynamic trafficking of geranylgeranylated-FBXO10 to the outer mitochondrial membrane compartment.

### Geranylgeranylated-FBXO10 is a client of PDE6δ-mediated transport to OMM

Our fluorescence recovery after photobleaching (FRAP) results suggested that an active molecular mechanism may be required to transport geranylgeranylated-FBXO10 through the aqueous cytosol to the OMM. We recently described PDE6δ, a prenyl-group binding protein that accommodates both farnesyl and geranylgeranyl isoprenoid moieties in its hydrophobic binding pocket, involved in the trafficking of geranylgeranylated-FBXL2 between the ER and PM. It is unknown if PDE6δ mediates the trafficking of prenyl-proteins to mitochondria, as no mitochondrial client(s) have been identified so far. Interestingly, PDE6δ co-purified with FLAG-FBXO10 during immunoaffinity purifications from enriched mitochondrial fractions, identified by unbiased mass spectrometry analysis (data available on request). We validated the mass spectrometric results using co-immunoprecipitations of endogenous PDE6δ with Strep-FBXO10 from whole cell lysates (Fig. 3A, top panel lane 2). F-box family members such as FBXO10 have leucine-rich repeats, WD repeats, and various other protein interaction domains involved in mediating binding to substrate and/or regulator proteins. We included nine more F-box family members in the co-immunoprecipitation assay to examine if PDE6δ is among the binders of this family. Of the ten F-box proteins tested, including FBXO11-the paralog of FBXO10, only one that bound endogenous PDE6δ possessed the C-terminal CaaX motif, i.e, FBXO10 (Fig. 3B and Fig. 1A). PDE6δ protein turnover was affected neither by CRISPR-Cas9 mediated-FBXO10 gene deletion nor by forced expression of FBXO10 (Fig. S3A, B), demonstrating that PDE6δ is not a substrate of FBXO10 for the ubiquitin-mediated degradation but rather a regulator client of FBXO10. Geranylgeranylation-deficient Strep-FBXO10(C953S) did not bind with PDE6δ (Fig. 3A, top panel lane 3), demonstrating the critical importance of the geranylgeranyl isoprenoid modification at C953 for the engagement with PDE6δ. Finally, deltarasin, a drug that inhibits the binding of prenyl-proteins by blocking the isoprenoid accommodating pocket of PDE6δ significantly inhibited the interaction of FBXO10 with PDE6δ (Fig. 3C). Our results demonstrate that PDE6δ interacts with FBXO10 *via* geranylgeranylation of its CaaX-motif cysteine953 and this interaction does not result in alterations of PDE6δ abundance. Thus, rather than a target for degradation, PDE6δ may serve as a regulator client of geranylgeranylated-FBXO10.

**Figure 3.**
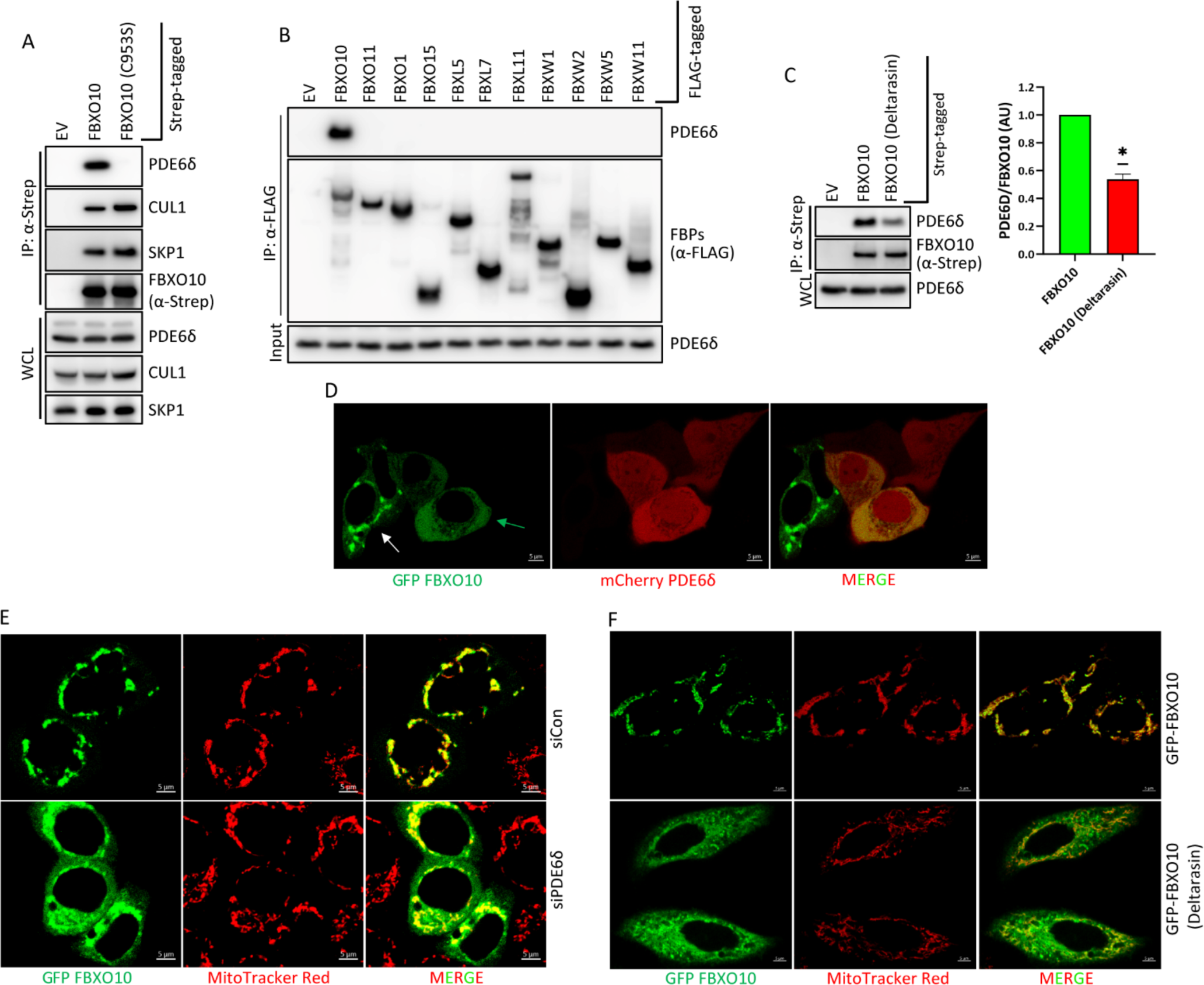
The lipid-binding PDE6δ chaperone mediates delivery of geranylgeranylated-FBXO10 to OMM **(A)** *PDE6δ binds FBXO10, but not geranylgeranylation-deEicient FBXO10(C953S).* STREP-tagged FBXO10, FBXO10(C953S) and empty vector (EV) were expressed in HEK293T cells. Post-cell harvesting STREP-Tactin immunoprecipitations were performed with standard protocols. Subsequently, immunoprecipitated complexes and corresponding whole cell lysates were subject to SDS-PAGE and immunoblotting as indicated. Representative of three independent experiments is shown. **(B)** *SpeciEic binding of PDE6δ to CaaX motif containing F-box protein, FBXO10.* FLAG-tagged indicated F-box family proteins representing FBXL, FBXW and FBXO subgroups and empty vector (EV) were expressed in HEK293T cells. Post cell harvesting anti-FLAG immunoprecipitations were performed with standard protocols. Subsequently, immunoprecipitated complexes were subject to SDS-PAGE and immunoblotting as indicated. Representative of two independent experiments is shown. **(C)** *Deltarasin treatment inhibits PDE6δ binding to FBXO10.* STREP-tagged FBXO10 expressing HEK293T cells were treated with deltarasin (2.5 µM) or DMSO overnight before cell harvesting. Deltarasin and DMSO treated frozen cell pellets were processed for STREP-Tactin immunoprecipitations with standard protocols. Subsequently, immunoprecipitated complexes were subject to SDS-PAGE and immunoblotting as indicated. Bar graph shows quantification of PDE6*δ* binding (n=2). Error Bar: SEM. Representative of two independent experiments is shown. **(D)** *PDE6δ depletion delocalizes FBXO10 away from OMM.* Non-targeting siRNA or siRNA targeting PDE6δ were transfected in HeLa cells for 20 hours as indicated. Post 20 hours, GFP-FBXO10 was expressed for 16 hours. Post treatment cells were treated with Mito-Tracker to decorate mitochondria before visualization by live cell confocal microscopy. Confocal images were captured using ZEIS LSM700 microscope. Shown are representative images of three independent experiments. Scale bar, 5 µm **(E)** *Changing the equilibrium between PDE6δ and FBXO10 re-distributes FBXO10 away from OMM.* GFP-FBXO10 and mCherry**-**PDE6δ were co-transfected in HeLa cells for overnight. Live cell confocal imaging of GFP-FBXO10 co-expressed with mCherry**-**tagged PDE6δ captured with ZEISS LSM750 confocal microscope. Green arrows indicate delocalization of GFP-FBXO10 in cells co-expressing mCherry-PDE6δ. white arrows indicate typical OMM distribution of FBXO10 in cells without mCherry-PDE6δ co-expressed. Shown is representative of three independent experiments. Scale bar 10µM. **(F)** *Deltarasin treatment delocalizes FBXO10 away from OMM.* Deltarasin (2.5µM) or DMSO treatment was carried out in HeLa expressing GFP-FBXO10 for 16 hours as indicated. Post treatment cells were treated with Mito-Tracker to decorate mitochondria before visualization by live cell confocal microscopy. Confocal images were captured using ZEIS LSM700 microscope. Shown is representative of two independent experiments. Scale bar: 5 µm.

PDE6δ enables active client prenyl-protein transport through the aqueous milieu of the cytosol between membrane compartments^14,26^. To examine the regulator function of PDE6δ for FBXO10, we tested the hypothesis that geranylgeranylated-FBXO10 is a client for PDE6δ mediated dynamic transport to the OMM. Extraction of client-prenyl proteins (e.g., FBXL2, NRAS and KRAS4B) from their steady-state membrane compartments upon forced expression of PDE6δ provides the assessment of PDE6δ function ^11,27^. We found FBXO10 extracted from mitochondria/OMM and rendered to cytosol upon overexpression of PDE6δ, as evident from a homogenous fluorescence of GFP-FBXO10 in cells co-transfected with mCherry-PDE6δ (Fig. 3D). Deltarasin, by blocking the interaction between PDE6δ and prenyl-protein (e.g., FBXL2, KRAS4B), paradoxically prevents prenyl-protein from associating with membrane compartments due to interruption in the regulated constant cycling between intended membrane compartments essential for steady-state subcellular localization^11,27^. We demonstrated that either the depletion of PDE6δ using siRNA silencing or the inhibition pharmacologically using deltarasin disrupts GFP-FBXO10’s localization to the mitochondrial membrane association (Fig. 3E, F and Fig. S3C, S3D) a result seen previously for FBXL2 and KRAS4B for their PM localization. Thus, we demonstrate that geranylgeranylated-FBXO10 is new mitochondrial client for PDE6δ, and our results, taken together, show PDE6δ is a cytosolic factor essential for the dynamic transport of geranylgeranylated-FBXO10 to the OMM.

### HSP90 and PDE6δ orchestrate the speciRic delivery of geranylgeranylated-FBXO10 to OMM

The primary structure of human FBXO10 polypeptide and its orthologs lack typical mitochondrial targeting signals (MTS). Although prenylation (farnesylation and geranylgeranylation) mediates the association with membrane compartments of diverse prenyl-proteins (small GTPase family, lamins), other prenyl-proteins do not show mitochondrial subcellular distribution (e.g., HRAS, KRAS4B, and FBXL2), unlike geranygeranylated-FBXO10 despite lacking MTS (Fig. 1B-D, Fig. 2A-G and Fig. 3D-F). Therefore, we sought to investigate the molecular determinant(s) necessary and sufficient for the specific delivery of geranylgeranylated-FBXO10 to the OMM. Client prenyl-proteins are delivered to PM and endomembrane by PDE6δ based on what is termed as “regulated second signal” usually juxtaposed to the CaaX sequences near C-termini of prenyl-clients (e.g., palmitoylation in case of HRAS, FBXL2, and polybasic sequence in the case of KRAS4B)^11,12^. In the absence of MTS and obvious second signal sequences in the primary structure of FBXO10 polypeptide, it is unclear how PDE6δ, although necessary (Fig. 3A-F), can deliver geranygeranylated-FBXO10 specifically to OMM. Cytosolic molecular chaperones HSP90 and HSP70 may mediate the mitochondria-specific protein targeting *via* docking onto integral OMM protein receptor TOM70^18^. Interestingly, by unbiased mass spectrometry analysis of enriched mitochondrial fractions, we found HSP90, HSP70, and TOM70 co-purified with FLAG-FBXO10 immunoaffinity purifications (data available on request). We validated the binding of endogenous HSP90, HSP70, and TOM70 with Strep-FBXO10 by their co-immunoprecipitation from whole cell lysates (Fig. 4A). Protein turnover of HSP90, HSP70, and TOM70 were affected neither by CRISPR-Cas9 mediated-FBXO10 gene deletion nor by forced expression of FBXO10 (Fig. S4A and Fig. S4B), showing that HSP90, HSP70, and TOM70 are not substrates of FBXO10 for the ubiquitin-mediated degradation but rather are regulator clients. Therefore, we examined the potential role of cytosolic HSP90 and HSP70 in the specific mitochondrial targeting of geranylgeranylated-FBXO10. We measured the mitochondrial redistribution of FBXO10 upon pharmacological intervention of HSP90 and HSP70 with CCT018159 and PES-Cl, respectively. Interestingly, our results show that inhibition of HSP90, but not HSP70, resulted in delocalization of FBXO10 away from mitochondria, as assessed by three independent assays, i.e., confocal immunofluorescence microscopy (Fig. 4B), flow cytometry of enriched mitochondrial fractions (Fig. 4C), and subcellular biochemical fractionations (Fig. 4D). Depletion of HSP90 and HSP70 with targeted-siRNAs further confirmed the redistribution of GFP-FBXO10 away from mitochondria upon reduction of HSP90 protein levels, but not HSP70 (Fig. 4E and Fig. S4C). Thus, our results reveal a predominant role for HSP90, but not HSP70, in specific mitochondrial targeting of FBXO10, as validated by the use of CCT018159 to demonstrate that the FBXO10 localization is dependent on the ATPase activity of HSP90. Furthermore, we found that whereas FBXO10 interacted with HSP90 and HSP70 and TOM70 (integral OMM receptor), geranylgeranylation-deficient FBXO10(C953S) failed to interact with TOM70 but maintained binding to HSP90 and HSP70 comparable to geranylgeranylated-FBXO10 (Fig. 4A, top panel lane 3). Loss of binding to TOM70, an integral OMM protein, is consistent with the subcellular re-distribution of geranylgeranylation-deficient FBXO10 (C953S) away from mitochondria (Fig 1A-D), suggesting that the absence of geranylgeranylation at C953 in C-terminal CaaX motif (Fig. 2D, E) and consequent lack of PDE6δ-mediated transport (Fig. 3A-E) thwarts stable mitochondrial association even in the presence of bound HSP90 and HSP70. Accordingly, CCT018159 treatment interrupted the mitochondrial localization of FBXO10 (Fig. 4B-D) despite no detectable impact on bound PDE6δ (Fig. 4F, lane 3 vs 2). Conversely, deltarasin treatment also interrupted the mitochondrial localization of FBXO10 (Fig 3E) without significant impact on bound HSP90 (Fig. 4F, lane 4 vs 2). Therefore, our results demonstrate that although both PDE6δ and HSP90 are necessary, individually neither is sufficient for maintaining the steady-state subcellular mitochondrial distribution of geranylgeranylated-FBXO10 (Fig. 3 and Fig. 4). These results suggest that there is a coordination between PDE6δ and HSP90 for the specific delivery of geranylgeranylated-FBXO10 to OMM and maintenance of steady-state OMM distribution. In support of the coordination between PDE6δ and HSP90 for the delivery of FBXO10 at OMM, we found PDE6δ co-immunoprecipitated with HSP90 and TOM70 in the presence of FBXO10, but not FBXO10(C953S) which is re-distributed away from mitochondria, demonstrating an OMM multi-protein complex of geranylgeranylated-FBXO10, aqueous trafficking factor PDE6δ, mitochondria specificity chaperone HSP90, and mitochondrial receptor TOM70 (Fig. 4G and Fig. S4D). Independent from the geranylgeranylation status of FBXO1O, we further assessed the impact of simultaneous inhibition of PDE6δ and HSP90 on the OMM re-distribution of GFP-FBXO10. Accordingly, we co-treated GFP-FBXO10 expressing cells with PDE6δ and HSP90 inhibitors (i.e., deltarasin and CCT018159) without inhibiting geranylgeranylation. CCT018159 (at half-dose) when treated simultaneously with deltarasin, resulted in efficient re-distribution of prenyl-FBXO10 away from OMM, assessed by FACS (Fig. 4H) and confocal imaging (Fig. S4E). Our results show coordinated roles of PDE6δ and HSP90 for the OMM distribution of prenyl-FBXO10, wherein PDE6δ enables transport of geranylgeranylated-FBXO10 through aqueous environment, and HSP90 directs FBXO10 docking on the mitochondrial receptor TOM70. Thus, PDE6δ and HSP90 together ensure the maintenance of steady-state subcellular OMM distribution and function of geranylgeranylated-FBXO10.

**Figure 4.**
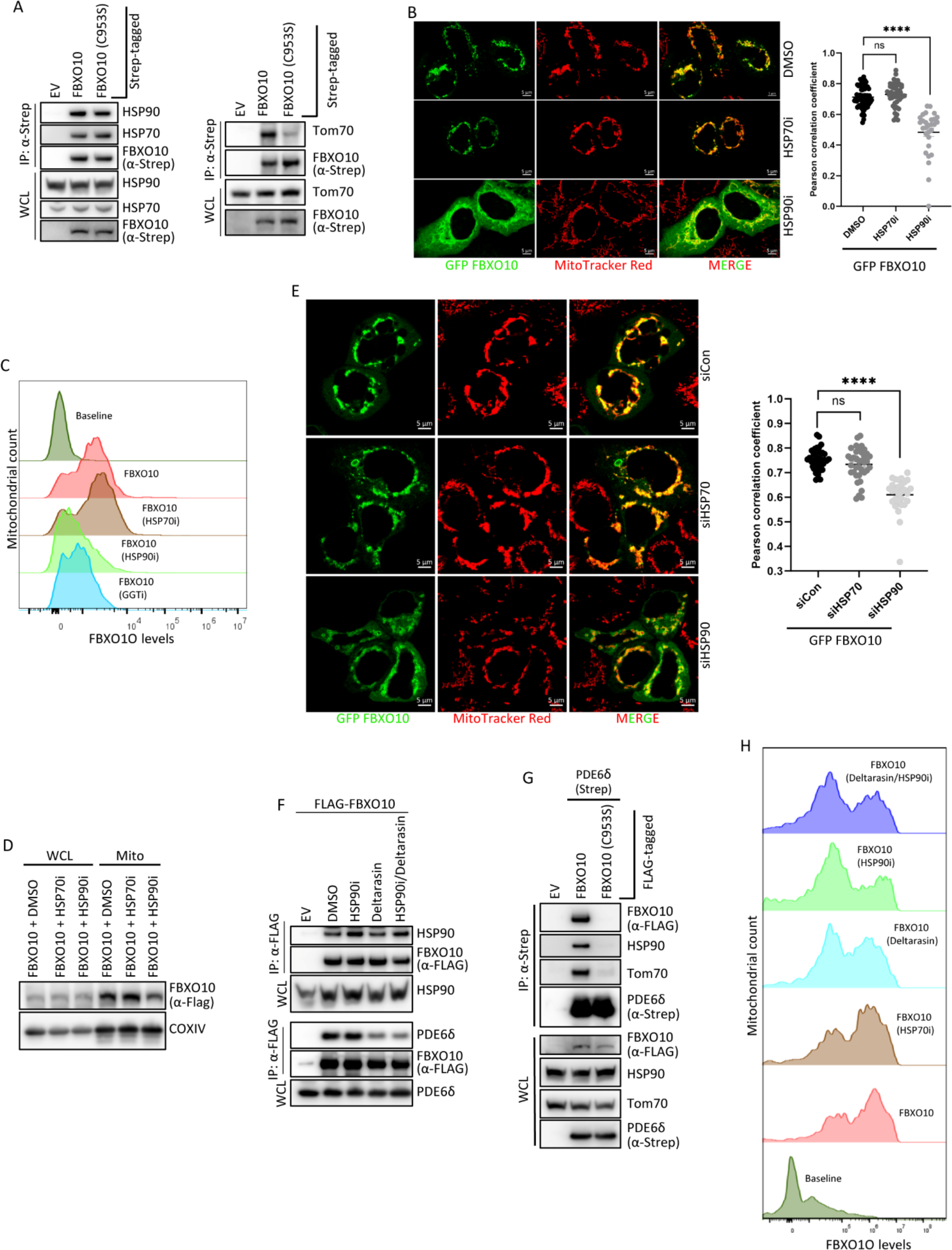
HSP90 coordinates with PDE6δ for the speciRic targeting of geranylgeranylated-FBXO10 to OMM **(A)** *HSP90 and HSP70 bind FBXO10 and FBXO10(C953S). TOM70 binds FBXO10 but not FBXO10(C953S).* STREP-tagged FBXO10 and FBXO10(C953S) were expressed in HEK293T cells before cell harvesting. Frozen cell pellets were processed for STREP-Tactin immunoprecipitations with standard protocols. Subsequently, immunoprecipitated complexes were subject to SDS-PAGE and immunoblotting as indicated. Shown is representative of three independent experiments. **(B)** *Inhibition of HSP90, but not HSP70, redistributes FBXO10 away from mitochondria.* CCT018159 (HSP90i: 10 µM), PES-Cl (HSP70i: 5 µM) or DMSO treatment was carried out in HeLa expressing GFP-FBXO10 for 16 hours as indicated. Post treatment cells were treated with Mito-Tracker to decorate mitochondria before visualization by live cell confocal microscopy. Confocal images were captured using ZEIS LSM700 microscope. Shown are representative images (n=70 for DMSO and n=47 for PES-Cl and n=29 for CCT018159). Pearson correlation coefficients (PCC) for the colocalization of GFP-FBXO10 and mitochondria were analyzed in different cells. Statistical analysis of plotted PPCs in scatter plots is show (right). p-values were calculated by student-T-test. Scale bar: 5 µm. **(C)** *Delocalization of mitochondria associated FBXO10 by inhibition of HSP90, but not HSP70.* Enriched intact mitochondrial fractions, as part of the experiment shown in Fig. 2C, were isolated from ptd-Tomato-FBXO1O expressing HEK293T cells treated with CCT018159 (10 µM), PES-Cl (10 µM) or DMSO for 16 hours by stepwise centrifugation. Before fractionation MitoView Green was added to non-expressing and ptd-Tomato-FBXO1O expressing cells undergoing treatments for 15 minutes to track mitochondria during flowcytometry analysis (FACS) to quantify mitochondria-associated ptd-Tomato-FBXO10 signal. Representative FACS histogram confirms FBXO10 at OMM (panel 2 and 3 from top) shown as rightward shifted peak in untreated and PES-Cl (HSP70i) treated samples from the baseline non-expressing controls (panel 1) and its loss from OMM due to CCT018159 (HSP90i) and GGTi treatments, but not PES-Cl, shown as leftward shifted peak closer/matching baseline non-expressing controls (panel 4, 5 from top). Representative of three independent experiments is shown. **(D)** *Loss of FBXO10 from mitochondrial fractions upon HSP90 inhibition.* HEK-293T cells expressing FLAG-tagged FBXO10 cells were treated with PES-Cl (10 µM), CCT018159 (10 µM) or vehicle DMSO for 16 hours. Post-harvesting cell lysates were subjected to subcellular fractionation using stepwise ultracentrifugation (see Methods). Enriched mitochondrial and cytosolic fractions were assayed by SDS-PAGE and immunoblotted as indicated. Representative of three independent experiments is shown. **(E)** *HSP90 and PDE6D binding to FBXO10 is mutually independent.* HEK293T cells expressing FLAG-tagged *FBXO10* were treated with deltarasin and/or CCT018159 overnight before cell harvesting. Frozen cell pellets were processed for STREP-Tactin immunoprecipitations with standard protocols. Subsequently, immunoprecipitated complexes were subject to SDS-PAGE and immunoblotting as indicated. Representative of two independent experiments is shown. **(F)** *HSP90, but not HSP70, silencing delocalizes FBXO10 away from OMM.* Non-targeting siRNA or siRNA targeting HSP90 and HSP70 were transfected in two consecutive cycles in HeLa cells for total of 40 hours. Post-20-hour silencing, GFP-FBXO10 was expressed for 20 hours. Post-treatments cells were treated with Mito-Tracker for ∼30 minutes to decorate mitochondria before visualization by live cell confocal microscopy. Confocal images were captured using ZEIS LSM700 microscope. Shown are representative images (N=38 for siRNA NT, N=37 siRNA *HSP70 and N=29 siRNA HSP90).* Pearson correlation coefficients (PCC) for the colocalization of GFP-FBXO10 and mitochondria were analyzed in different cells. Statistical analysis of plotted PPCs in scatter plots is shown (right). p-values were calculated by student-T-test. Scale bar: 5 µm. **(G)** *PDE6δ co-immunoprecipitated mitochondrial FBXO10 and TOM70.* STREP-tagged *PDE6δ* was co-expressed with FLAG-FBXO10 and FLAG-FBXO10(C953S) in HEK293T cells before cell harvesting. Frozen cell pellets were processed for Strep-Tactin immunoprecipitations with standard protocols. Subsequently, immunoprecipitated complexes were subject to SDS-PAGE and immunoblotting as indicated. Representative of two independent experiments is shown. **(H)** *Simultaneous inhibition of HSP90 and PDE6D efEiciently delocalizes FBXO10 away from OMM.* Enriched intact mitochondrial fractions were isolated from ptd-Tomato-FBXO1O expressing HEK293T cells treated with DMSO, CCT018159, deltarasin and in combination as indicated for 16 hours by stepwise centrifugation. Post fractionation MitoView Green was added to non-expressing and ptd-Tomato-FBXO1O expressing cells undergoing treatments for ∼30 minutes to track mitochondria during flowcytometry analysis (FACS) to quantify mitochondria-associated ptd-Tomato-FBXO10 signal. Representative FACS histogram confirms FBXO10 at OMM (panel 2 from bottom) shown as rightward shifted peak from the baseline non-expressing controls (panel 1) and its loss from OMM due to treatments with CCT018159 (HSP90i) alone (panel 5), deltarasin alone (panel 4), and combination of deltarasin and CCT018159 (panel 6), shown as leftward shifted peaks closer/matching baseline non-expressing controls.

### Geranylgeranylation-deRicient FBXO10(C953S) impairs mitochondria-driven myogenic differentiation in iPSCs and murine myoblasts

Muscle dysfunctions in patients on statins (widely used clinical drugs for lowering blood cholesterol) forces suspension of statin therapy^25^. Statins inhibit the cellular mevalonate pathway, which otherwise produces geranylgeranyl isoprenoids. Interestingly, our results demonstrate that lovastatin treatment inhibits FBXO10 trafficking to mitochondria, which depends on its geranylgeranylation at C953, (Fig. 2A). Skeletal muscles (SM) are highly enriched in mitochondria, and, as such, SM are critically dependent on efficient mitochondrial function. A balance between mitochondrial fission and fusion is critical for myogenic differentiation^28,29^, however, mitochondria-driven myogenesis and underlying molecular mechanism(s) are not well understood. Although several studies have suggested dysfunctional mitochondria impair myogenic differentiation, the major focus has centered on long-term nucleus-driven genome remodeling and transcriptional reprogramming during myogenesis. How mitochondria-driven local actions are activated swiftly in response to myogenic cues to maintain optimal mitochondrial network plasticity and functional efficiency is not well understood. We uncovered a new geranylgeranylated-E3 ligase, FBXO10, dynamically distributed at OMM (Fig. 1A-D, Fig. 3 and Fig. 4), and found its re-distribution away from OMM results in dysregulation of OMM proteostasis (Fig. 1F, G and Fig. S1F-H), impairment of mitochondrial morphological dynamics (Fig. IK), impaired mitochondrial-ATP production (Fig. 1H, I) and mitochondrial-membrane potential (Fig. 1J) and a potential downstream molecular target of statins known to cause muscle dysfunctions in patients (Fig. 2A). Therefore, we examined if the maintenance of dynamic OMM subcellular distribution of geranylgeranylated-FBXO10 impacts mitochondria-driven myogenesis. To test this hypothesis, we first expressed geranylgeranylation-deficient FBXO10(C953S) in human iPSCs and murine myoblasts (C2C12 cells). We generated human iPSCs stably expressing FBXO10, geranylgeranylation-deficient FBXO10(C953S) and vehicle lentiviral infected controls (EV). Quantitative RT-PCR analysis showed comparable FBXO10 and FBXO10(C953S) expressions in engineered iPSCs (Fig. S5A). Moreover, we confirmed that there was no detectable impact on pluripotency potential, as assessed by OCT4 immunostaining among the EV, FBXO10 and FBXO10(C953S) expressing iPSCs (Fig. S5B). We subjected engineered and controlled iPSCs to a 20-day myogenic differentiation program and monitored the impact sequentially at progenitors, myoblasts, and final skeletal muscle myotubes. At the progenitor iPSC stage (Fig. S5B) and up to the myoblast stage as assessed by MyoD immunostaining (Fig. S5C), no detectable differences were observed between the controls (EV) and engineered iPSCs, suggesting no obvious differentiation defects up to the myoblast stage. Interestingly, we found remarkable differences at the final skeletal muscle myotube stage as assessed by immunostaining of myotube marker, myosin heavy chain (MyH) (Fig. 5A). Whereas myotubes (MyH positive) were visualized as denser and longer networks upon FBXO10 expression (Fig. 5A, middle vs top panels), in contrast expression of geranylgeranylation-deficient FBXO10(C953S) completely blocked formation of normal myotubes (Fig. 5A, bottom panel). Since the myogenic differentiation defect due to expression of geranylgeranylation-deficient FBXO10(C953S) was remarkably obvious onwards from myoblast stage (iPSC derived model), we carried out a series of experiments in C2C12 murine myoblasts which are known to recapitulate the human myogenic program ^30,31^ and are pliable to genetic and pharmacologic interventions compared to human iPSCs.

**Figure 5.**
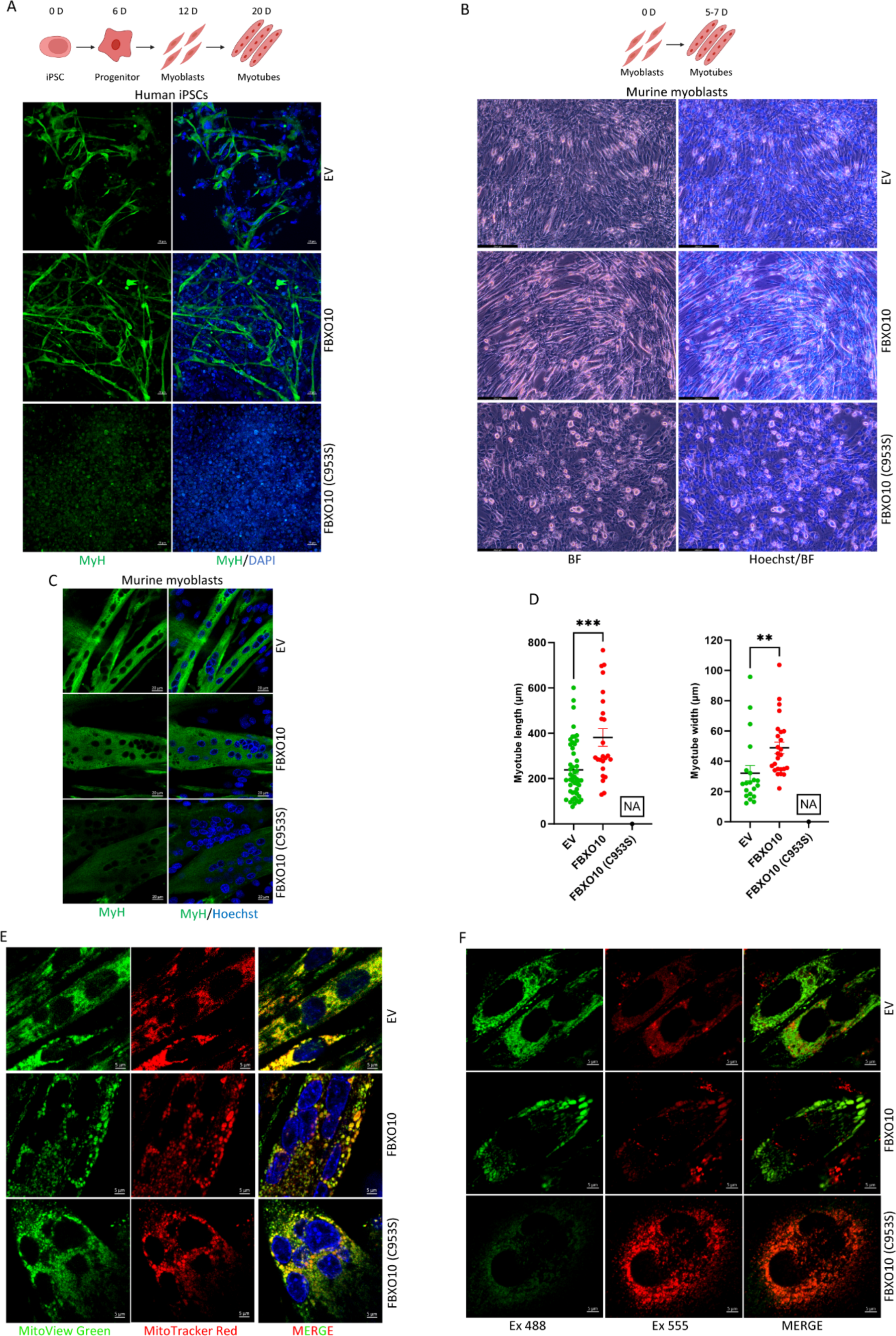
Geranylgeranylation-deRicient FBXO10(C953S) impairs mitochondria-driven myogenic differentiation in iPSCs and murine myoblasts. **(A)** *Geranylgeranylation-deEicient FBXO10(C953S) impairs myotube formation in iPSCs.* Human iPSCs stably expressing FLAG-FBXO10, FLAG-FBXO10(C953S) and vehicle controls (EV) were differentiated for 20 days to myotubes (see protocol details in methods section). At the end point, the samples were fixed and immunostained with myosin heave chain (MyH) antibody conjugated to A488 fluorescent probe. Hoechst was added to stain nuclei prior to visualization by ZEISS LSM700 confocal microscopy. Scale bar: 20 µm. **(B)** *Geranylgeranylation-deEicient FBXO10(C953S) blocks myotube formation in murine myoblasts C2C12.* Murine C2C12 myoblasts stably expressing FLAG-FBXO10, FLAG-FBXO10(C953S) and vehicle controls (EV) were differentiated for 5 days to myotubes (see protocol details in methods section). At the end point, the samples were treated with Hoechst, and visualized by microscopy by live cell brightfield and florescence microscopy. Scale bar: 235.6 µm. **(C)** *Myosin heavy chain positive myotube formation is blocked by geranylgeranylation-deEicient FBXO10(C953S).* Murine C2C12 myoblasts stably expressing FLAG-FBXO10, FLAG-FBXO10(C953S) and vehicle controls (EV) were differentiated for 7 days to myotubes (see protocol details in methods section). At the end point, the samples were fixed and immunostained with myosin heave chain (MyH) antibody conjugated to A488 fluorescent probe. Hoechst was added to stain nuclei prior to visualization by ZEISS LSM700 confocal microscopy. Scale bar: 20 µm. **(D)** *Geranylgeranyl-FBXO10, but not FBXO10(C953S), promotes myotube size increase.* Indicated differentiated samples processed as in 5C were analyzed for myotube size measurements. Lengths and widths of MyH positive myotubes in EV and FBXO10 expressing samples were calculated using imageJ software and scatter plots were generated using GrapPad Prism software. FBXO10 (C953S) expressing samples were not analyzed (NA) because of non-detectable MyH positive myotubes (Myotube length: n=49 for EV and n=24 for FBXO10; Myotube width: n=20 for EV and n=25 for FBXO10). **(E)** *Geranylgeranylation-deEicient FBXO10(C953S) promotes abnormal myotube formation and mitochondrial damage.* Murine C2C12 myoblasts stably expressing FLAG-FBXO10, FLAG-FBXO10(C953S) and vehicle controls (EV) were differentiated for 8 days for myotube formation. At the end point samples were treated with MitoTraker Red, MitoView Green, and Hoechst to decorate mitochondria and nuclei. Representative images, captured by by ZEISS LSM700 confocal microscopy, show mitochondrial and myotube morphology. (Scale bar, 5 µm). **(F)** *Persistent mitophagy promoted by geranylgeranylation-deEicient FBXO10(C953S).* Mitophagy probe Mito-Kiema was stably co-expressed in murine C2C12 myoblasts expressing FLAG-FBXO10, FLAG-FBXO10(C953S) and vehicle controls (EV). As indicated, samples were differentiated for 7 days for myotube formation. At the end point samples were analyzed for mitophagy status by live cell confocal microscopy using 488 and 555 excitation lasers. Representative images, captured by ZEISS LSM700 confocal microscopy show mitochondria (green) and mitochondria/lysosomal(red) compartments. Red: indicates mitophagy positive. Green: indicates mitophagy negative.

The analysis of myogenic differentiation using C2C12 myoblasts revealed remarkable defects, highly similar to those seen in the iPSC model, in the derived myotubes due to expression of geranylgeranylation-deficient FBXO10(C953S) expressed at levels comparable to that of FBXO10 (Fig. 5B, Figs. S5D and S5E). Whereas FBXO10 promoted formation of oversized myotubes, geranylgeranylation-deficient FBXO10(C953S) expression resulted in the formation of abnormal myotubes visualized as round and multinucleated (nuclei either clustered and/or malorganized) and inhibited myogenic differentiation (Fig. 5B and Fig. S5E). Myotube size measurements, visualized by MyH immunostaining, show that FBXO10-expressing myotubes are longer and broader (perhaps fused sideways) compared to EV controls (Fig. 5C, D). In contrast, insignificant MyH immunostaining is observed in abnormal myotubes expressing geranylgeranylation-deficient FBXO10(C953S), precluding their size measurements. Taken together, analysis of both iPSCs and C2C12 derived myotubes show that geranylgeranylation-deficient FBXO10(C953S) expression counteracts execution of proper myogenic differentiation program, correlating with defective mitochondrial proteostasis, morphological dynamics, and energetics (Fig. 1).

To examine if the defective myogenic differentiation of C2C12 myotubes caused by the expression of FBXO10 (C953S) resulted from dysregulated mitochondrial morphological dynamics and/or mitochondrial damage, we examined mitochondrial network dynamics in live C2C12 cells with MitoTracker Red (stains intact healthy mitochondria) and mitochondrial mass measuring dye MitoView Green (stains both healthy and damaged mitochondria). First, we found during the differentiation program from myoblasts through myotubes that the mitochondrial morphology was different and notable (Fig. S5F). Whereas mitochondria in myoblasts appear like a typical reticulate network spread across the myoblasts in sparse and shorter sub-networks, in contrast the mitochondria in myotubes appear fused and are organized in dense and long networks (Fig. S5F, top vs bottom). Interestingly, whereas we visualized a typical myotube mitochondrial morphological profile of dense and long networks in myotubes expressing empty vector, the FBXO10 expressing myotubes contained hyperfused, round, and larger mitochondria appearing clustered around perinuclear regions (Fig. 5E, middle vs top panels). In contrast, the abnormal myotubes due to geranylgeranylation-deficient FBXO10 (C953S) expression contained strikingly altered mitochondrial morphologies, visualized as small, fragmented, and irregularly organized (Fig. 5E, bottom panel). Moreover, the analysis for the incorporation of MitoTracker Red (a marker for healthy, intact mitochondria) and MitoView Green (a marker for total mitochondria mass) revealed that FBXO10-expressing myotubes contained mostly healthy intact mitochondrial networks assessed by the maximum overlap between markers (Fig. 5E, middle panels). However, in contrast, the overlap between markers was limited in abnormal myotubes expressing geranylgeranylation-deficient FBXO10 (C953S), with significant portions stained only green, revealing either damaged or fragmented mitochondrial networks (Fig. 5E, bottom panels). Consistent with mitochondrial damage and fragmentation, the mitochondrial-membrane potential was inhibited by the expression of geranylgeranylation-deficient FBXO10 (C953S) (Fig. 1J and Fig. S1L). Prompted by induction of mitochondrial damage and fragmentation, we analyzed if geranylgeranylation-deficient FBXO10 (C953S) impacted the mitophagy of damaged mitochondrial networks. To measure mitophagy *in vivo*, we used mitochondria targeted mt-Keima based imaging assay (a fluorescent probe for measuring mitophagy *in vivo* based on pH sensitivity)^32^ and analyzed C2C12 myotubes upon expression of wild type FBXO10, geranylgeranylation-deficient FBXO10 (C953S), and vehicle controls (EV). We found mitophagy in FBXO10-expressing myotubes was similar to what we observed in EV controls (Fig. 5F, top vs middle panels); in contrast, geranylgeranylation-deficient FBXO10 (C953S)-expressing abnormal myotubes were undergoing enhanced mitophagy (Fig. 5F, bottom panels) consistent with their damaged and fragmented mitochondrial networks (Fig. 5E and Fig. 1J, 1K).

### FBXO10 loss impairs myogenic differentiation and PGAM5 degradation at OMM during myogenic differentiation

To determine if FBXO10 is required for myogenic differentiation, we first performed FBXO10 silencing with two different shRNAs directed at coding sequence (shCDS) and 3’ untranslating region (sh3’UTR) and analyzed the impact on C2C12 myoblasts to differentiate into skeletal muscle myotubes. Both shRNAs (shCDS and sh3’UTR) directed a significant silencing of FBXO10 expression (Fig. S1M) and resulted in a defective myogenic differentiation compared to non-targeting shRNA control, assessed by immunostaining of C2C12 myotubes with MyH (Fig. 6A). Whereas typical and organized myotubes were visualized in non-silenced controls, in contrast shCDS and sh3’UTR silencing resulted in a striking reduction of intact/organized myotubes (Fig. 6A panels 1-3, and bargraph) and MyH-stained abnormal myotubes appeared sharply reduced and malorganized (Fig. 6A, top vs 2^nd^ and 3^rd^ panels from top). Interestingly, exogenous supplementation with FBXO10 rescued intact/organized myotube formation in sh3’UTR silenced myoblasts, significantly although incompletely. (Fig. 6A, bottom panel, bar graph and Fig. S6A). Moreover, mitochondrial ATP production was completely rescued to control levels in sh3’UTR silenced myotubes supplemented with exogenous FBXO10 (Fig. 6B). Thus, both silencing of FBXO10 and expression of geranylgeranylation-deficient FBXO10 (C953S) impair myotube formation consistent with their similar impacts on mitochondrial morphological dynamics, fragmentation, and damage (Fig. 5, Fig. 6A, Fig. 1J, 1K and Fig S1M). Next, we analyzed mitophagy using mt-Keima assay after FBXO10 silencing in C2C12 abnormal myotubes. Our results show a typical mitochondrial morphological network profile, dense and long, in control C2C12 myotubes with basal/limited mitophagic mitochondrial networks (Fig. 6C top panel, and Fig. S1M). However, in shCDS and sh3’UTR silenced myotubes, mitochondrial networks clustered in perinuclear regions and show a marked increase in mitophagy consistent with their mitochondrial damage and fragmentation (Fig. 6C and Fig. S1M).

**Figure 6.**
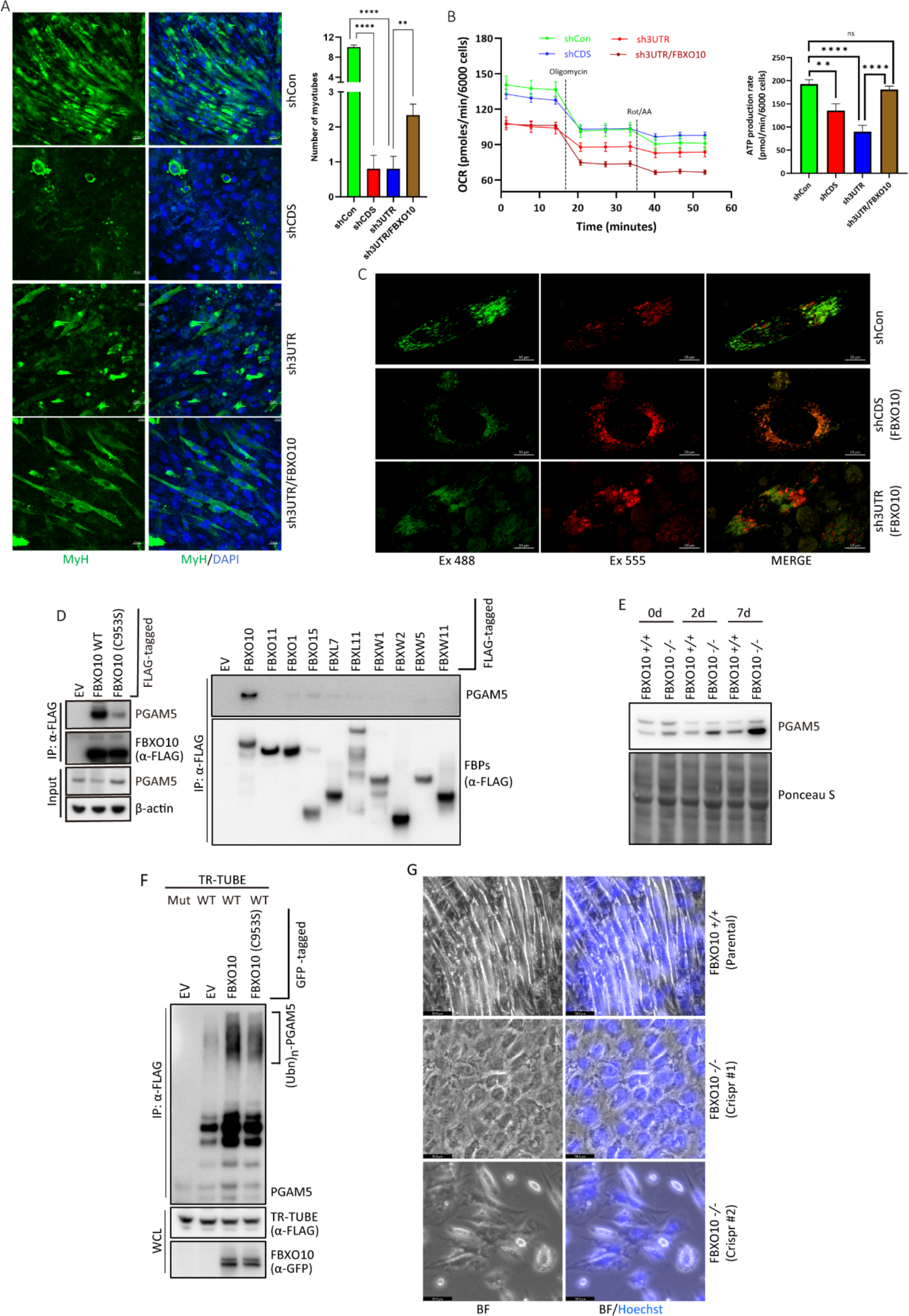
FBXO10 loss impairs myogenic differentiation. PGAM5 is targeted for FBXO10 mediated degradation during myogenic differentiation. **(A)** *FBXO10 reconstitution rescues defective myotube formation.* FBXO10 depletion was carried out with two independent shRNAs targeted to untranslated (3’UTR) and coding (CDS) regions in C2C12 myoblasts. In parallel, FLAG-FBXO10 expression was reconstituted in 3’UTR shRNA treated C2C12 myoblasts (sh3’UTR/FBXO10), as described in methods section. Control shRNA, FBXO10 depleted, and FBXO10 reconstituted C2C12 myoblasts were subjected to myogenic differentiation for 7 days to generate myotubes. At the end point, the samples were fixed and immunostained with myosin heave chain (MyH) antibody conjugated to A488 fluorescent probe to stain myotubes. Hoechst was added to stain nuclei prior to visualization by ZEISS LSM700 confocal microscopy. Number of properly formed myotubes was counted in indicated samples (N=7 control shRNA, N=10 shCDS, N=10 3UTR and N=15 sh3’UTR/FBXO10, where N represents the fields of view). Quantification of myotube formation is shown in the bargraph. p-values were calculated by student-T-test. Scale bar: 20 µm. **(B)** *FBXO10 reconstitution rescues defective mitochondrial ATP production.* FBXO10 depletion and reconstitution was carried out as in (A). Control shRNA, FBXO10 depleted, and FBXO10 reconstituted C2C12 myoblasts were subjected to myogenic differentiation for 7 days days. Measurement of mitochondrial ATP production was carried out by Seahorse XF Real-Time ATP Assay. Line graphs show oxygen consumption rate (OCR), and bar graph shows mitochondrial and glycolytic ATP production rate obtained from OCR and ECAR measurements. Bar graph represent quantification from 8 biological replicates. Error bars: SEM. **(C)** *FBXO10 depleted myotubes show persistent mitophagy.* Mito-Kiema, a mitochondria-specific mitophagy probe, was stably co-expressed in murine C2C12 myoblasts expressing control-shRNA, two independent shRNAs targeted to untranslated (3’UTR) and coding (CDS). As indicated, samples were differentiated for 7 days for myotube formation. At the end point samples were analyzed for mitophagy as in Fig. 5G. Red: indicates mitophagy positive. Green: indicates mitophagy negative. **(D)** *PGAM5 speciEically interacts with FBXO10.* Left: C2C12 myoblasts stably expressing FLAG-FBXO10, FLAG-FBXO10(C953S) and vehicle control (EV) were differentiated for 2 days. Post-2-day differentiation, samples were harvested and processed for whole cell lysate immunoprecipitation with ani-FLAG-resin. Immunoprecipitated complexes and WCLs were subject to SDS-PAGE and immunoblotting as indicated Shown is representative of three independent experiments. Right: FLAG-tagged FBXO10, indicated F-box family proteins and empty vector (EV) were expressed in HEK293T cells. Post cell harvesting anti-FLAG immunoprecipitations were performed with standard protocols. Subsequently, immunoprecipitated complexes were subject to SDS-PAGE and immunoblotting as indicated. Shown is representative of two independent experiments. **(E)** *PGAM5 levels Eluctuate during myogenic differentiation.* Myogenic differentiation in Parental C2C12 and CIRISPR/CAS9 deleted FBXO10 myoblast clone was carried out for up to 7 days before sample harvesting at indicated time points (see methods sections for details). Frozen cell pellets were lysed, and whole cell lysates were processed for SDS-PAGE followed by immunoblotting as indicated. **(F)** *FBXO10, but not FBXO10, promotes PGAM5 polyubiquitylation.* FLAG-tagged TR-TUBES, wild type and mutant, were expressed in HeLa cells as indicated. Immunoprecipitations from the prepared whole cell lysates were carried out with anti-FLAG resin. The immunopurified samples were subjected to SDS-FGAE followed by immunoblotting with the indicated antibodies. The ladder of bands corresponding to polyubiquitylated PGAM5 is marked by the bracket on the right. Immunoblot of TR-TUBEs expression is shown at the bottom. Shown is the representative of two independent experiments. **(G)** *Crispr/Cas9-mediated FBXO10 deletion blocks myogenic differentiation.* Parental C2C12 and two independent CIRISPR/CAS9 deleted FBXO10 clones (CRISPR#1 and CRISPR#2), as indicated, were differentiated for 7 days. At the endpoint, the samples were treated with Hoechst, and visualized by brightfield and immunofluorescence microscopy (Scale bar, 59.9 µm).

Among the 18 OMM putative substrates of FBXO10 identified by the comparative mass spectrometry screen (Fig. 1F-G), PGAM5 (a mitochondrial phosphatase) is implicated for the regulation of stem cell differentiation, mitochondrial morphological dynamics, and mitophagy^33–36^, i.e., phenotypes impacted by FBXO10 depletion and/or expression of geranylgeranylation-deficient FBXO10 (C953S). We confirmed PGAM5 indeed is a mitochondrion-associated protein by co-localization with a mitochondrial marker, cytochrome c, using confocal immunofluorescence imaging (Fig. S6B). We found that treatment with proteasome inhibitor MG132 and/or CUL-neddylation inhibitor MLN4924 increased the PGAM5 protein levels in differentiating C2C12 myoblasts (Fig. S6C). Based on these results, we examined if PGAM5 is targeted for degradation *via* FBXO10 during myogenic differentiation program and conversely if deregulated degradation of PGAM5 may explain the defective differentiation seen by FBXO10 depletion or expression of geranylgeranylation-deficient FBXO10 (C953S). First, by unbiased mass spectrometry analysis of FBXO10 immunocomplexes, purified from enriched mitochondria, we found PGAM5 specific peptide enrichment (data available on request). To validate our mass spectrometry data, we confirmed indeed FBXO10 binds endogenous PGAM5 (Fig. 6D). The binding of PGAM5 to FBXO10 is dependent on OMM distribution, as geranylgeranylation-deficient FBXO10 (C953S), which distributes away from OMM, shows significant loss in PGAM5 binding (Fig. 6D and Fig. S6D). Accordingly, we found FBXO10, but not FBXO10(C953S), promoted PGAM5 decrease in differentiating C2C12 myoblasts (Fig. 6D, input). PGAM5 specifically engages FBXO10, as tested for 10 F-box proteins including its paralog FBXO11 (Fig. 6D). Next, we examined the impact on PGAM5 degradation and myogenic differentiation upon FBXO10 gene deletion *via* CRISPR/CAS9 editing in C2C12 myoblasts (Fig. S6E). Our results show PGAM5 protein levels fluctuate during myogenic differentiation, with noticeable turnover during the myogenic differentiation program (Fig. 6E and Fig. S6F). However, FBXO10 deletion prevented PGAM5 protein degradation during C2C12 differentiation program (Fig. 6E). Moreover, FBXO10 promoted polyubiquitylation of endogenous PGAM5 at OMM whereas basal PGAM5 polyubiquitylation and polyubiquitylation upon expression of geranylgeranylation-deficient FBXO10 (C953S) were comparable as assayed by ubiquitin-binding TR-TUBE assay. (Fig. 6F). Finally, we examined two independent FBXO10 knock out myoblast clones (CRISPR#1 and CRISPR#2) during myogenic differentiation to myotubes. Our results show both myoblast clones are strikingly defective in skeletal muscle myotube formation visualized by brightfield and immunofluorescence microscopy (Fig. 6G). Moreover, MyH staining confirmed defective myogenic differentiation, abnormal skeletal muscle myotube formation and reduction in intact/organized myotubes in both FBXO10 knock out clones consistent with shRNA-mediated FBXO10 silencing (Fig. S6G and Fig. 6A).

Taken together, our results show re-distribution of FBXO10 away from mitochondria and FBXO10 loss (silencing and/or deletion) result in an analogous impact on mitochondrial morphological dynamics, mitochondrial ATP production, mitochondrial membrane potential, and mitophagy during myogenic differentiation. We show PGAM5 is polyubiquitylated and targeted for degradation *via* FBXO10 at OMM during myogenic differentiation program. Thus, we expose geranylgeranylated-FBXO10 as a new OMM resident regulator of mitochondria-driven biological processes such as myogenic differentiation program

## Discussion

Cellular membranes harbor nearly a quarter of the total human proteome, and a majority of current therapeutics (>60%) target membrane proteins. Protein turnover of membrane-associated proteins at specific subcellular membrane compartments *via* E3-ubiquitin ligases is not well understood. For example, the steady-state subcellular distribution of most E3-ligases and the underlying mechanisms to maintain their spatial profile are unknown. A comprehensive understanding of how E3 ligases are spatially restricted to accomplish the compartment specific protein turnover is warranted. The emergence of E3-ligase mediated targeted protein degradation strategies, such as PROTACs/molecule glues for therapeutic purposes ^42–44^, highlights the need for understanding such mechanisms because it would pave the way for compartment specific therapeutic targeting of proteins. FBXO10, a substrate receptor of a CRL1, was uncovered by our studies as dynamically trafficked E3 selectively to the outer mitochondrial membrane (OMM) (Fig. 3, Fig. 11-D). Additionally, we demonstrated the mechanism for its specific delivery and maintenance at the OMM. Pertinently, we gained insights into functional consequences by the re-distribution of FBXO10 away from OMM, as an example to underscore the importance of the spatially restricted ubiquitylation process. FBXO10, is deregulated in lymphomas and also implicated various biological processes ^37–41^. As inspection of FBXO10 primary structure revealed an atypical CaaX motif, *CTIL*, conserved across orthologs, we examined whether FBXO10 is a candidate for prenyl isoprenoid modification (i.e., either 15-carbon farnesyl or 20-carbon geranylgeranyl), to regulate its OMM association. Our data shows indeed that FBXO10 is modified by geranylgeranyl isoprenoid (Fig. 2D, E). The acceptor site for geranylgeranylation is cysteine953 (C953) located in the C-terminal CaaX motif of FBXO10 (Fig. 1A, B and Fig. 2D, E). By swapping the C953 to S953 to preclude isoprenoid modification in FBXO10(C953S) mutant we demonstrated that the integrity of C953 is indispensable for the OMM distribution of geranylgeranylated FBXO10 (Fig. 1A-D, Fig. S1C and Fig. 4G). Geranylgeranylation-deficient FBXO10(C953S) re-distributed away from the OMM to the cytosol, as well as nuclear compartments (Fig. 1A-D, Fig. S1C); a profile similar to soluble non-prenylated proteins as has been recognized in the case of small GTPase RAS proteins, HRAS, NRAS, and KRAS ^45,46^, suggesting that FBXO10 substrates are membrane associated. Whereas geranylgeranylated FBXO10 at OMM assembled with its CRL subunits CUL-1 and SKP1, so did the soluble FBXO10(C953S) (Fig. 1E). However, reduced neddylated-cullin-1 associated with soluble FBXO10(C953S) (Fig. 1E). Neddylation/deneddylation cycle of CRLs is regulated by substrate engagement^22,23^. Thus, the redistribution away from OMM precludes FBXO10(C953S) from the substrate engagement at OMM.

We exploited geranylgeranylation-deficient FBXO10(C953S) for the identification of potential substrates of FBXO10 at OMM by label-free-quantitative mass spectrometry (LFQ-MS/MS) of enriched mitochondrial preparations isolated from cells expressing FBXO10 and FBXO10(C953S). One of the top predicted OMM candidates for selective degradation *via* FBXO10, identified by unbiased LFQ-MS/MS, included PGAM5. PGAM5 is a mitochondrial phosphatase, which was independently validated as a substrate at OMM (Fig. 6F-H), underscoring the fidelity of this approach to identify compartment specific substates for the spatially restricted E3s. Functionally, our studies suggest that the timed PGAM5 degradation *via* FBXO10 promotes myogenic differentiation program by maintaining homeostasis of efficient and healthy mitochondrial networks. It is known that PGAM5 activity, in part, promotes the balance between mitophagy and compensatory mitochondrial biogenesis during stem cell differentiation^36^. PGAM5 activity promotes the mitophagic clearance of damaged or fragmented mitochondria, ensuring constant rewiring and biogenesis of only healthy mitochondrial networks during stem cell differentiation. We propose that FBXO10-mediated degradation of PGAM5 may serve to dampen and/or terminate non-selective mitophagy, enabling proper progression through a differentiation program. Interestingly, PGAM5, when in the cytosol, is a target of KEAP1, a substrate receptor of a CRL3, and oxidative stress counteracts KEAP1-PGAM5 interaction to promote mitophagy ^47,48^. Since KEAP1 is predominantly a cytoplasmic E3, it is not entirely clear if PGAM5 is degraded in the cytosol before its targeting to mitochondria or post-proteolytic processing at mitochondria. FBXO10 targets PGMA5 only at OMM as demonstrated by the geranylgeranylation-deficient FBXO10(C953S), which is re-distributed away from OMM, losses PGAM5 binding, and inhibits PGAM5 degradation (Fig. 6E and Fig. S6D). Nonetheless, it may be argued that cytosolic KEAP1 regulates events before the initiation of mitophagy program and FBXO10 regulates events at the OMM to terminate mitophagy, for example, as during myogenic differentiation. Thus, the FBXO10-mediated degradation of PGAM5 at OMM represents an acute and selective mitochondria-driven degradation event to maintain a rapid check of non-selective mitophagy. Consistent with this notion, PGAM5 proteins levels fluctuate, in part dependent on FBXO10 mediated degradation, during myogenic differentiation program (Fig. 6F). Increased PGAM5 levels, either by FBXO10 loss or expression of FBXO10(C953S), correlates with mitochondrial damage and persistent mitophagy, which counteracts appropriate progression through myogenic differentiation program (Figs. 5 and 6). Although we show PGAM5 degradation *via* FBXO10 during mitochondria-driven myogenic differentiation, there are likely additional physiologic functions of FBXO10 due to other substrates at the OMM (data available on request). For example, AKAP1/MIRO1 regulation (data available on request) may impact mitochondrial morphological dynamics and clustering, while hexokinase regulation (data available on request) can impact metabolism, perhaps also important for the differentiation process. Taken together, our work unveils FBXO10 as a critical local OMM resident E3 ubiquitin ligase to swiftly respond upon myogenic cues enabling faithful execution of myogenic program.

Our studies of impairment of mitochondrial trafficking of FBXO10 may explain key aspects of human muscle dysfunction. For example, statins are widely used drugs in clinics to lower blood cholesterol. Mechanistically, statins inhibit HMG-CoA reductase, a key step in the cellular mevalonate pathway also involved in the geranylgeranyl isoprenoid synthesis. Statins reduce cardiovascular mortality and morbidity, but one of the most common causes for patients discontinuing statin therapy is the occurrence of muscle weakness and cramps referred to as statin-associated muscle symptoms (SAMS)^25^. Our studies thus identify geranylgeranylated-FBXO10 as a potential molecular target that can help explain statin-toxicity in muscles. We suggest statins prevent prenylation and therefore trafficking of FBXO10 to mitochondria (Fig. 2A), thus impairing mitochondrial homeostasis and muscle differentiation, as shown by our iPSC derived and murine myoblast myogenic differentiation models. Moreover, it is interesting to note that the isoprenoid geranylgeranyl pyrophosphate which is required for the lipidation of FBXO10 is generated *via* the mevalonate pathway in the endoplasmic reticulum but, the starting substrate of the mevalonate pathway is acetyl-CoA, a key metabolite generated by mitochondria. Thus, a possibility exists that the geranylgeranyl pyrophosphate levels may itself form part of a regulatory loop of mitochondrial proteostasis involving acetyl-CoA levels which, in turn, regulate the extent of FBXO10 localization to mitochondria.

We believe that geranylgeranylation itself, although essential for membrane association, is insufficient to dictate OMM compartment specificity. For example, extensively documented prenyl-GTPases (HRAS, NRAS, KRAS) and prenyl-FBXL2 are delivered to PM due to “second signals” involving palmitoylation sites and/or poly basic sequence in conjunction with CaaX sequence prenylation^11,12^. Additionally, the lack of classical MTS signal, and no obvious “second signal” sequence in FBXO10 primary structure, promoted us to investigate the molecular determinants necessary and sufficient for the trafficking and maintenance of dynamic OMM distribution of geranylgeranylated FBXO10. We show that the combined actions of two cytosolic factors, PDE6δ and HSP90, orchestrate the specificity of prenyl-FBXO10 towards OMM distribution (Fig. 3 and Fig. 4). Both cytosolic factors, PDE6δ and HSP90, are necessary but neither is sufficient to accomplish the OMM trafficking of FBXO10. PDE6δ, by shielding of the hydrophobic geranylgeranyl modification at the CaaX motif enables FBXO10 trafficking in the aqueous cellular environment, whereas the chaperone HSP90 specifically directs FBXO10 to the OMM receptor TOM70. Thus, these studies uncover a previously unknown function of PDE6δ to direct trafficking to mitochondria and identify prenyl-FBXO10 as the first OMM client for the PDE6δ. Moreover, we show how a prenyl-protein without an inherently coded “second signal” in the primary structure, recruits a second cytosolic factor HSP90, thus resulting in OMM-specific trafficking. There may be additional proteins which are similarly trafficked to the OMM despite lacking a mitochondrial targeting sequence *via* the tag-teaming of PDE6δ and HSP90, and this opens the possibilities for other discoveries in future.

Functionally, we show that the OMM distribution of geranylgeranylated-FBXO10 is critical for mitochondrial homeostasis (Fig. S7), whereas lack of FBXO10 OMM localization or FBXO10 depletion results in impaired mitochondrial dynamics, reduced mitochondrial ATP production, and inhibited mitochondrial membrane potential. Several E3-ligases, such as Parkin or FBXL4^19,20^, are known to translocate to the OMM, but they typically trigger mitophagy and wholesale mitochondrial recycling. The precision targeting of selective OMM proteins such as PGAM5 by FBXO10 allows for more fine-tuned responses in which the cellular machinery selectively degrades specific mitochondrial proteins without triggering mitophagy. In this context, FBXO10 spatial distribution at OMM is not static but rather dynamic, as demonstrated by two independent assays. First, excess PDE6δ partitioned FBXO10 (Fig. 4D) and second, FBXO10 levels recovered at OMM following focal photobleaching of GFP-FBXO10 (Fig. 2H). Thus, understanding regulation of ON and OFF recruitment of FBXO10 at OMM may help gain insights into the events involved in programmed degradation of various OMM resident substrates. We posit that FBXO10 may gain access to specific OMM targets, in part, dependent on cues regulating FBXO10 traffic to and from OMM. Decoding specific cues, for example, that promote and/or counteract PDE6δ and HSP90 binding to prenyl-FBXO10 will require follow up studies. In summary, our findings propose a model of selective mitochondrial proteostasis *via* FBXO10 (Fig. S7) which paves the way for understanding physiological and pathophysiological roles of E3-ligase mediated selective OMM protein degradation complementing known E3-ligase pathways which regulate mitophagy.

## Acknowledgements

Authors acknowledge Dr. Said Sebti for sharing GGTi-2418 and Drs. Jalees Rehman and Michele Pagano for inputs for the manuscript preparation. This work was supported by the grant from the National Institutes of Health R35 GM137452 (to S.K.) and funds provided by the BCMG department at UIC (to S.K.)

## Author contributions

SAB: Conceptualization, performed most of the biochemical, cell genetic, myogenic differentiation (iPSCs and murine C2C12 myoblasts), autoradiography, mitophagy, FACS, fractionations and imaging experiments, analyses of proteomic data, writing-original draft, literature survey, and prepared figures.

ZV: Performed biochemical, cell genetic, imaging experiments and statistical analysis of the data along with SAB, prepared figures with SAB.

LJ: Imaging, FRAP, myogenic differentiation (C2C12 myoblasts) and biochemical experiments, participated in figure preparation and manuscript writing.

SS: Developed assay and computational analysis to validate CRISPR-Cas9 deleted FBXO10 clones, confirmed shRNA and overexpression clones by real time qPCR and prepared the model figure.

RF: Performed CRISPR-Cas9 mediated FBXO10 deletion in C2C12 myoblasts, and biochemical assay along with SAB.

AG: Cloned mouse FBXO10 CRISPR oligos. Developed FBXO10(*ΔF-box*) mutant and biochemical assay along with SAB.

HI: Performed immunoblotting assays during myogenic differentiation. RA: Participated in autoradiography with SAB.

UB and AD: Performed mass spectrometry analysis of samples prepared by SAB.

SK: Supervision, conceptualization, design, interpretation, manuscript writing, review, and editing, project administration, funding acquisition.

All authors discussed the results and commented on the manuscript.

## Declaration of interests

None.

## Inclusion and diversity statement

We support inclusive, diverse, and equitable conduct of research.

## Reagents and Resources

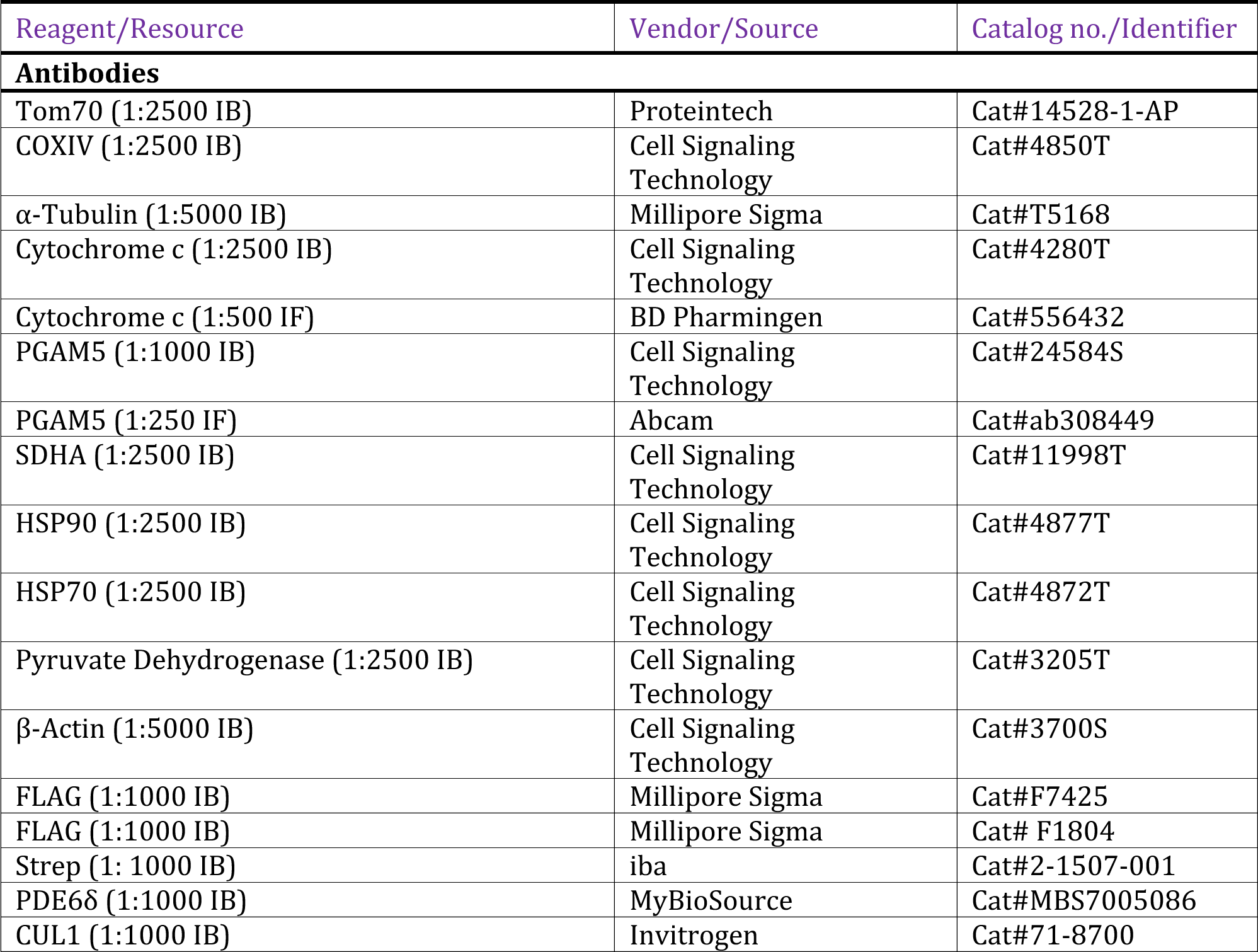
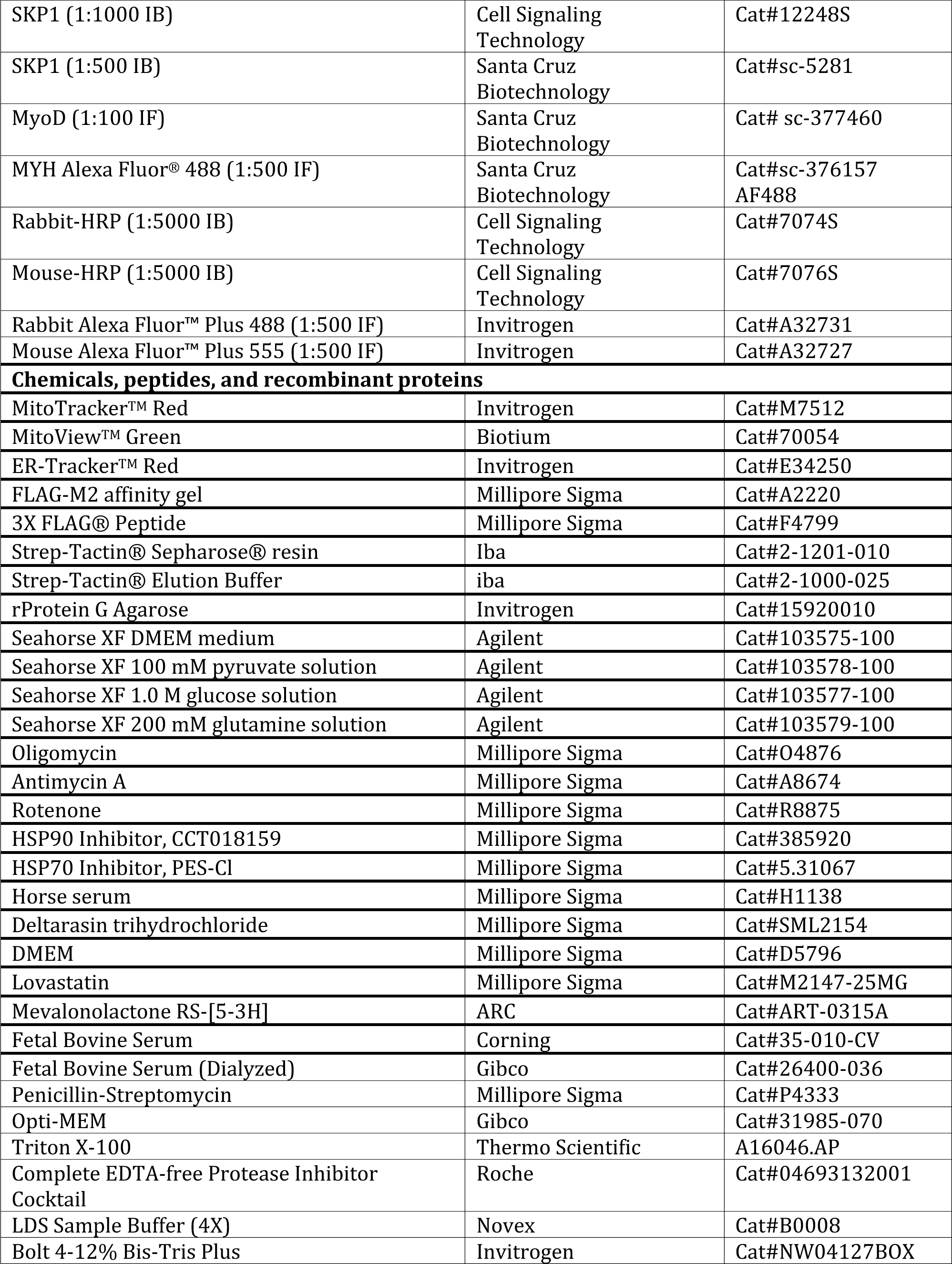
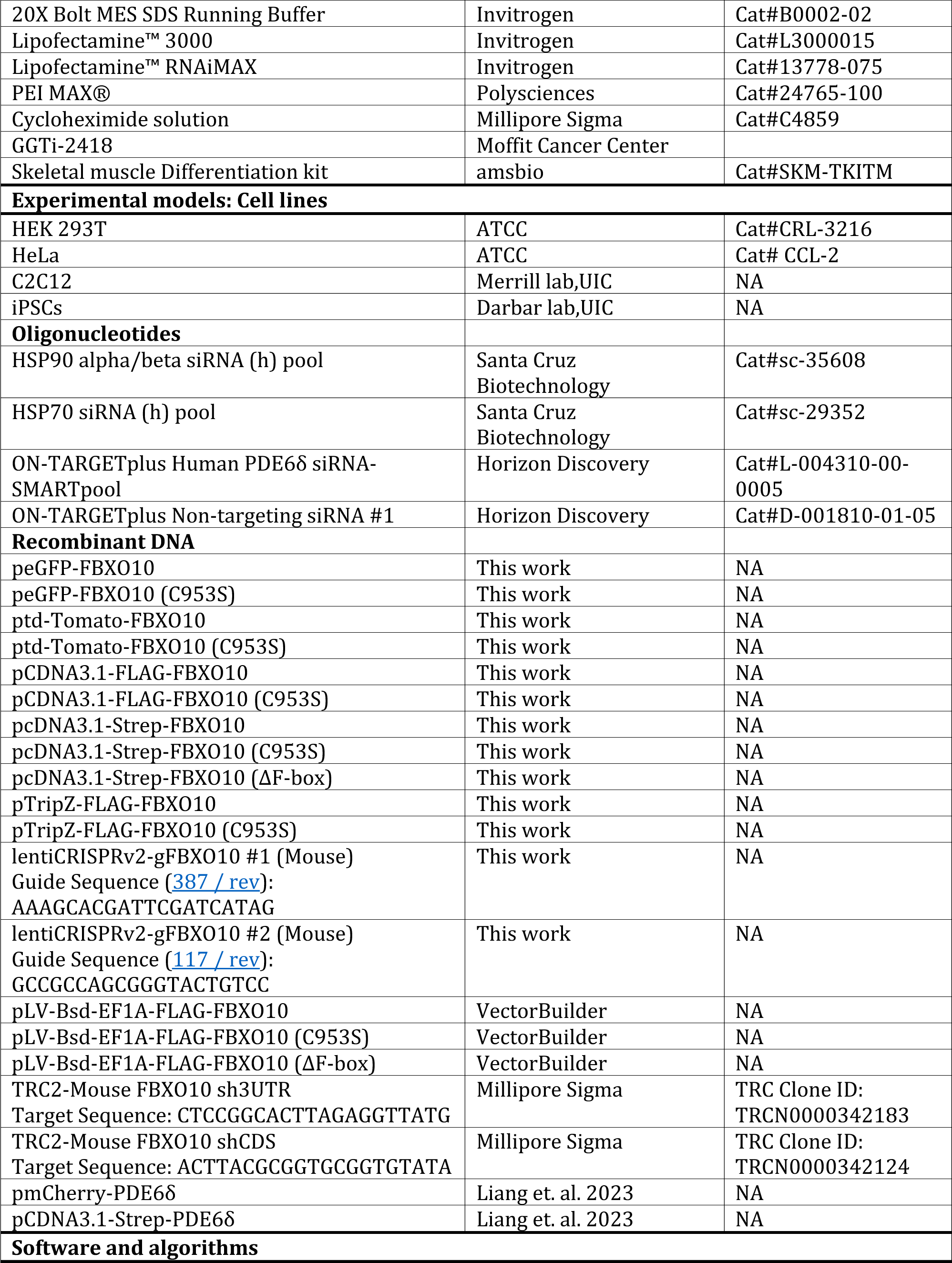
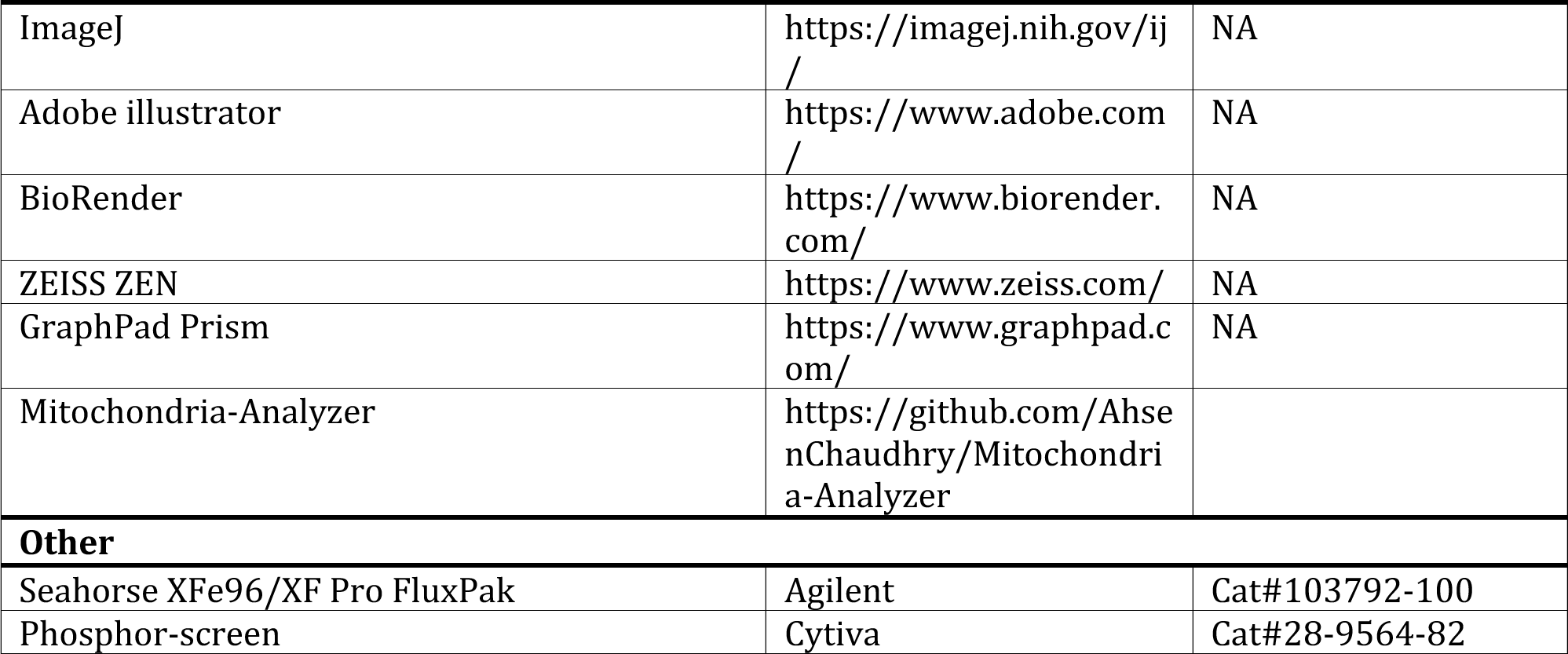

## Methods

### Metabolic labeling and prenylation

H^3^-mevalonolactone based prenylation assay was performed as described previously^49–51^. HEK293T stable cell lines expressing FBXO10 and FBXO10 (C953S) were grown in 6 cm dishes and treated with 15 μM lovastatin for about 2 hrs. 2 mL of labeling medium (DMEM containing 5% dialyzed serum, 0.2 mCi/mL of concentrated H^3^-mevalonolactone and 15 μM lovastatin) was added to the dishes. After labeling overnight, cells were lysed, and FLAG tagged proteins were immunoprecipitated using anti-FLAG M2 affinity gel (Sigma). Elution was carried out with 2X Bolt LDS sample buffer and the samples were blotted onto the PVDF membrane. The membrane was placed under a phosphor screen (Cytiva) in a cassette and incubated at room temperature in dark. The screen was exposed in a phosphor imager (Amersham Typhoon, GE) for [H^3^] detection. The PVDF membrane was finally immunoblotted for the expression of FLAG-tagged FBXO10 and FBXO10 (C953S).

### Label-free-quantiRication mass spectrometry

Enriched mitochondria were isolated from HEK293T cells stably expressing FBXO10 and FBXO10 (C953S) and vehicle control (EV) by stepwise centrifugation. Samples were lysed in SDS based lysis buffer (50 mM Tris Ph7.4, 150 mM NaCl, 1% SDS, 1mM EDTA and 50mM NaF) containing protease inhibitors (Roche) and sonicated. Protein lysates were processed using S-Trap (Protifi) following manufacturers protocol. In brief, samples were reduced with 20 mM DTT for 1 hour at 57°C, alkylated with 50 mM iodoactamide for 45 minutes at room temperature in the dark and subsequently the reaction quenched with another aliquot of 20 mM DTT. The reduced and alkylated samples were acidified with 12% phosphoric acid. Samples were loaded onto the S-trap (Protifi) and washed three times with S-trap binding buffer and subsequently digested with 500 ng Trypsin at 47°C for 1h. Peptides were eluted using a solution of 40% acetonitrile followed by 80% acetonitrile in 0.5% acetic acid. The organic solvent was removed using a SpeedVac concentrator and the samples were reconstituted in 0.5% acetic acid. Peptides were loaded onto an Acclaim PepMap trap column (2 cm x 75 µm) in line with an EASY-Spray analytical column (50 cm x 75 µm ID PepMap C18, 2 μm bead size) using the autosampler of an EASY-nLC 1000 HPLC (Thermo Scientific). Solvent A was 2% acetonitrile in 0.5% acetic acid and solvent B was 80% acetonitrile in 0.5% acetic acid. The gradient was held for 5 min at 5% solvent B, ramped in 60 min to 15% solvent B, in 35 min to 25% solvent B, in 20 min to 40% solvent B, and in 10 min to 100% solvent B. The peptides were gradient eluted directly into a Thermo Scientific Orbitrap Eclipse Mass Spectrometer. High resolution full spectra were acquired with a resolution of 240,000, an AGC target of 1e6, with a maximum ion time of 50 ms, and a scan range of 400 to 1500 m/z. All precursors with charge states between 2-7 were considered for fragmentation. All MS/MS HCD spectra were collected in the ion trap using the rapid scan mode, with an AGC target of 2e4, maximum inject on time of 18 ms, one microscan, 0.7 m/z isolation window, auto scan range mode, and normalized collision energy (NCE) of 27. The instrument was set to run at top speed with a cycle time of 3 seconds. Dynamic exclusion was set to 30 seconds. The MS/MS spectra were searched against the Uniprot human reference proteome using the Andromeda search engine^52^ integrated into the MaxQuant environment using the following settings: oxidized methionine (M), and deamidation (asparagine and glutamine) were selected as variable modifications, and carbamidomethyl I as fixed modifications; precursor and fragment mass tolerance was set to 10 ppm. The identifications were filtered using a false-discovery rate (FDR) of <1% at the protein and peptide level using a target-decoy approach. Only proteins with at least two unique peptides were reported. Data analysis was performed using Perseus ^53^. The protein intensities were log 2 transformed and normalized using the median. The results were further filtered for proteins that are identified in three replicates of at least one group. Missing values were imputed from a normal distribution. To identify proteins differing in intensity between the three groups, a Student’s t-test was applied and corrected for multiple hypothesis testing using Benjamini-Hochberg ^54^. A total of 4215 proteins were quantified across all 9 samples. Proteins with a fold change of ≥2 (log 2 FBXO10/FBXO10 (C953S) ≥1 and ≤-1) and an FDR of ≤5 % were considered ‘deregulated’ and Functional Annotation Clustering analysis was performed on deregulated proteins in FBXO10 vs FBXO10 (C953S) using DAVID software (https://david-d.ncifcrf.gov/)^55^. Terms with significant enrichment scores (ES ≥ 2, *P* ≤ 0.01) were plotted in a graph. The mass spectrometric raw files are accessible at https://massive.ucsd.edu under accession MassIVE MSV000094240 and at www.proteomexchange.org under accession PXD050382.

### Subcellular fractionation

Subcellular fractionation of cells was performed as described previously^56^. Briefly, HEK293T cells were harvested and resuspended in the fractionation buffer (225 mM mannitol, 75mM sucrose, 0.1 mM EGTA, 30 mM Tris pH 7.4) containing protease inhibitors. Samples were kept on ice for 20 min and subsequently processed by Dounce homogenization. Cell debris was separated by centrifuging the samples 2 times at 600 × g for 5 min. Crude mitochondria were isolated at 7000 × g for 10 min and washed once with the fractionation buffer. The supernatant was centrifuged at 100,000 × g for 1 h to separate the light membranes (pellet) and the cytosol (supernatant). The required fractions were processed for downstream applications.

### Protease protection assay

Cells were harvested and mitochondria were isolated as described previously in the methods section. Mitochondria were resuspended in the fractionation buffer without protease inhibitors and treated with trypsin (50μg/mL) for 15 min-1 h on ice. Post trypsin treatment, mitochondria were pelleted down at 7000 × g for 10 min. For western blotting mitochondria were lysed and the lysates were immunoblotted for the expression of FBXO10 and various mitochondrial proteins. For flow cytometry, mitochondria were treated with MitoView Green in the fractionation buffer prior to trypsin treatment. Post trypsin treatment, mitochondria were washed once with the fractionation buffer to remove the residual trypsin, resuspended in the fractionation buffer, and analyzed using CytoFLEX S flow cytometer (Beckman Coulter) and FlowJo software.

### ATP rate assay

XF Real Time ATP Rate Assay was performed using the Agilent Seahorse XFe96 Extracellular Flux Analyzer according to the manufacturer’s instructions. C2C12 cells were plated in Agilent Seahorse 96-well XF Cell Culture Microplate. Injection ports A and B were loaded with 10X Oligomycin (10µM) and 10X Rotenone/Antimycin A (5µM each) respectively. The data was analyzed using the Wave software.

### Myogenic differentiation

*C2C12 myoblasts to myotubes:* For differentiation, growth medium of C2C12 myoblasts was changed to differentiation medium (DM: DMEM containing 2% Horse Serum and 1× Pen-Strep). DM was changed every 1-2 days for 5-7 days. Myotubes were obtained at this stage.

*iPSCs to myotubes:* iPSCs were differentiated to Skeletal muscle myotubes using SKM-KITM - Skeletal muscle Differentiation kit (amsbio) according to manufacturer’s guidelines. Briefly, iPSCs were plated onto collagen I coated plates and grown in Skeletal Muscle Induction Medium for 6 days with a medium change after every 2 days. Myogenic precursors formed were dissociated using 0.05% Trypsin-EDTA and plated onto collagen I coated plates at 5000 cells/cm^2^. Cells were grown in Skeletal Myoblast Medium for 6 days with media change after every 2 days. Myoblasts were observed at this stage. Medium was switched to Myotube Medium for 8 days with a medium change every 3 days. Myotubes were obtained at this stage.

*Cell culture:* All cell lines were grown at 37°C and 5% CO2 concentration in a humidified incubator. HEK293T, HeLa and C2C12 cells were maintained in DMEM supplemented with 10% FBS and 1× Pen-Strep. iPSCs were grown on vitronectin coated plates containing PluriSTEM® Human ES/iPS Cell Medium (Sigma).

### CRISPR-Cas9 gene editing, gene silencing and molecular biology

*CRISPR/Cas9 deletion:* Guide oligos for FBXO10 knockdown were designed using the CRISPOR program. LentiCRISPRv2 vector which also expresses Cas9 was used for cloning guide oligos using standard protocols. Lentiviral transductions were carried out to stably express the guide RNA and Cas9. Puromycin was used for selecting clones with genomic integration. PCR amplification from genomic DNA encompassing the PAM site followed by sequencing was used to confirm FBXO10 knockout clones.

*Gene silencing and viral transduction:* HeLa cells were transfected with ON-TARGETplus siRNAs (Dharmacon) and Santa Cruz Biotechnology siRNAs using Lipofectamine RNAiMAX (Invitrogen) according to the manufacturer’s instructions. siRNAs were transfected on two consecutive days. Overexpression transfections were done on the second day post second siRNA transfections using Lipofectamine 3000. Silencing was confirmed by immunoblotting using antibodies against the silenced proteins. Transfections and viral transductions were carried out using standard procedures. Transfections of HEK293T and HeLa cells were performed using PEI MAX® and Lipofectamine 3000 respectively. For retroviral transductions, Gag-Pol and VSVG were used for virus packaging. For Lentiviral infections, pΔ8.2 and VSVG were used for the packaging.

*Cloning and site directed mutagenesis:* GFP, ptd-Tomato, FLAG and Strep tagged FBXO10 in pCDNA3.1 vector; Strep tagged FBXO10 (ΔF-box) in pCDNA3.1 vector; and FLAG tagged FBXO10 in pTRIPZ lentiviral vector were cloned using standard molecular cloning procedures. GFP, ptd-Tomato, FLAG and Strep tagged FBXO10 (C953S) mutants in pCDNA3.1 were developed using QuikChange Site-Directed Mutagenesis procedure. Flag tagged FBXO10 (C953S) mutant in pTRIPZ was made by digesting out the C-terminal region of FBXO10 in pTRIPZ and swapping with PCR amplified FBXO10 (C953S) mutant digested with same enzymes. Confirmation of mutation or deletion was done by sequencing.

*Real time quantitative PCR:* RNA was isolated from harvested cells using RNeasy Mini Kit (Qiagen). cDNA synthesis was conducted using RNA to cDNA EcoDry™ Premix (Oligo dT) (Takara). Real-time quantitative PCR (RT-qPCR) was performed using SYBR green in a CFX Duet Real-time PCR machine (Bio-Rad). Gene expression was quantified by (ΔΔ*Cq*) method using GAPDH was as the normalization control.

### Microscopy and Rluorescence recovery after photobleaching (FRAP)

*Live cell and immunoEluorescence microscopy:* For confocal microscopy studies, HeLa and C2C12 cells were plated on glass bottom dishes. Transient transfections, if needed, were done using Lipofectamine 3000 reagent. For experiments requiring differentiation, C2C12 myoblasts were differentiated on glass-like polymer bottom dishes. GFP and mcherry tagged proteins, specific membrane trackers and Hoechst nuclear dye were used for live cell microscopy. For immunofluorescence studies, cells were fixed and incubated with either fluorescent dye tagged primary antibody or primary antibody followed by fluorescent dye tagged secondary antibody. Nuclei were stained using Hoechst or DAPI stains. Imaging was performed using Zeiss LSM-700 confocal microscope and images were processed using Zeiss Zen software. For non-confocal live cell microscopy studies, C2C12 cells were grown and differentiated in 6 well plates. Nuclei were stained with Hoechst stain. Imaging was performed using Leica DMi8 microscope and images were processed using Leica LAS X software.

*FRAP:* Fluorescence recovery after photobleaching experiments were performed using a ZEISS LSM700 confocal microscope coupled with a temperature-, humidity- and CO2-controlled top-stage incubator for live-cell imaging. Images were acquired using a 63x oil immersion objective (Carl Zeiss). Three regions of interest (ROIs) were defined: ROI-1 was the indicated circular region; ROI-2 was un-photobleached region of similar size; and ROI-3 was defined as background and drawn outside of the cell, where its signal was subtracted from those of ROI-1 and ROI-2. Photobleaching was carried out with 488 nm laser at 100% power for 20 iterations. Images were acquired before and immediately after photobleaching every 1s for 90s. The laser power before and after photobleaching was kept identical. Fluorescence intensity of the ROI (F) at each time point was measured using imageJ software. FRAP curves (F(t)) were calculated: F(t)=100×(F(t)_FRAP region-_ F(t)_background)_/ (F_prebleach,FRAP region_- F(t)_background_). The data are normalized to the background corrected prebleach intensity (F_prebleach,FRAP region_) and multiplied by 100 to yield a percentage of recovery. The normalized data was used to calculate recovery rates: (Recovery rate=F_(reaching plateau for 1st time after photobleaching)_-F_(bleaching)_)/area/duration of reaching plateau for 1st time after photobleaching). Data were processed as above and plotted using Prism.

### Flow cytometry

HEK293T cells were transfected with ptd-Tomato FBXO10 and treated with various inhibitors as required. Mitochondria were isolated as described previously in the methods section and treated with Mitoview Green in the fractionation buffer for gating the mitochondria. FBXO10 peak and shifts due to various treatments were analyzed using CytoFLEX S flow cytometer (BECKMAN COULTER) and FlowJo software.

### Cycloheximide chase, immunoprecipitation, and immunoblotting

Transfected HEK293T cells were harvested and lysed in lysis buffer (50 mM Tris pH 7.4, 150 mM NaCl, 1% TritonX_100_, 50 mM NaF, 1 mM EDTA, 50 mM NaF) containing protease inhibitors. The samples were sonicated (pulse on: 10 s, pulse off: 10 s, and Amplitude: 30%) in Sonic Dismembrator with Cup Horn (Fisher Scientific). Lysates were pre-cleared using Protein G Agarose (Invitrogen) by rocking for 1–2 h at 4°C (16 rpm). Immunoprecipitions were done using anti-FLAG M2 (Sigma) or Strep-Tactin Sepharose 1h-overnight and the samples were eluted using 3X-FLAG peptide or Strep-Tactin elution buffer respectively. Samples were denatured in Bolt LDS sample buffer at 70°C for 10 min and processed for immunoblotting using standard protocols. For cycloheximide chase, HEK293T cells were transfected with EV and FBXO10 expressing constructs. Approximately 2 days post transfection, cycloheximide was added at a concentration of 100 μg/ml for different time periods. Cells were lysed and subjected to immunoblotting. C2C12 cells were differentiated for 2 days for myogenic program prior to cycloheximide treatment.

### QuantiRication and statistical analysis

Pearson’s correlation coefficients for colocalization were calculated using the Zeiss Zen software per single cell. Western blot quantification was carried out by densitometric analysis using Image Studio Lite software (LI-COR Biosciences). Mitochondrial morphology was analyzed using Mitochondria Analyzer Image J/Fiji plugin. Statistical analysis was performed using GraphPad Prism software. Unpaired Student’s t-test was performed when comparing two groups. Significance was defined as ns = p > 0.05, * = p ≤ 0.05, ** = p ≤ 0.01, *** = p ≤ 0.001, **** = p ≤ 0.0001. Error bars represent the standard error of the mean (SEM).

## Notes

### Competing Interest Statement

The authors have declared no competing interest.

