## Supplemental Figures for "Geranylgeranylated-SCF^FBXO10^ Regulates Selective Outer Mitochondrial Membrane Proteostasis and Function"

**Supplementary Figures**

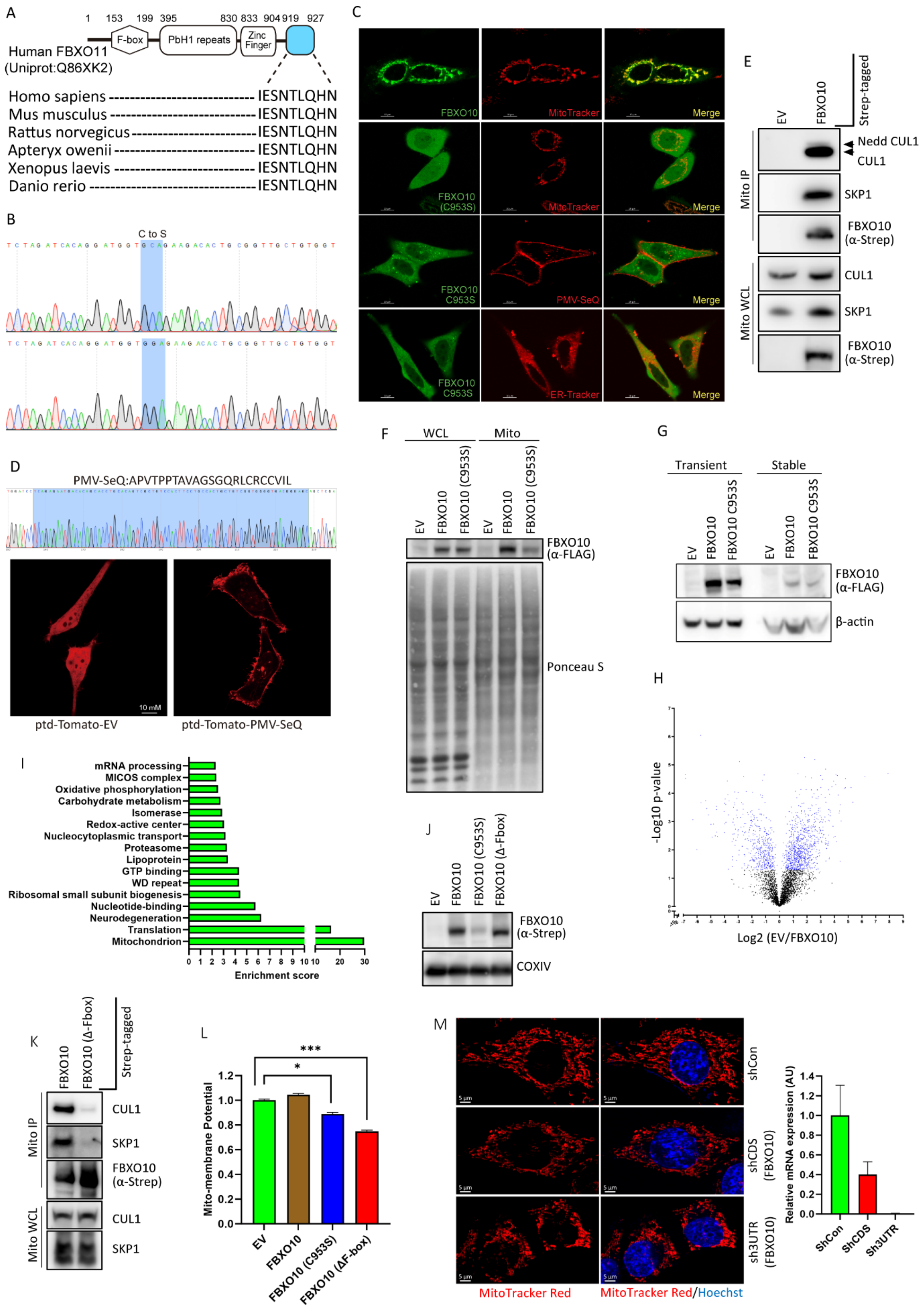


Bhat *et al.*, Figure S1

**Figure S1. FBXO10 distributed at mitochondria regulates mitochondrial homeostasis.**

**(A)** *FBXO11 polypeptide, unlike FBXO10, does not terminate in a CaaX motif.* FBXO11 schematic at top shows the predicted protein domain configuration highlighting F-box, PBH1. The alignment of C-terminal amino acid sequences of FBXO11 orthologs is shown.

**(B)** *CaaX-motif mutant of FBXO10 generated by site-directed mutagenesis.* The Sequence alignment of wild type cDNA and CaaX-mutant cDNA generated by site-directed mutagenesis shows the codon swap for the Cysteine953 to Serine953.

**(C)** *FBXO10 subcellular distribution at mitochondrial networks.* HeLa cells expressing GFP-FBXO10 (panel 1) and CaaX-motif mutant GFP-FBXO10(C953S) (panels 2, 3 and 4) were either treated with Mito-Tracker, ER-Tracker or co-transfected with PMV-SeQ to decorate mitochondrial networks (panels 1 and 2), ER (panel 4) and plasma membranes (PM, panel 3), respectively. ER-Tracker and Mito-Tracker were added 15 minutes prior to live cell confocal imaging and analysis as in Fig. 1B.

**(D)** *PMV-SeQ is a newly generated florescent probe for plasma membrane and motile vesicles*. HeLa cells were transfected by either pTd-tomato vector or pTd-tomato-PMV-SeQ cDNA constructs and subjected to confocal imaging captured by ZEISS-LSM700 microscope. The pTd-tomato-PMV-SeQ fluorescent-probe is designed based on juxtaposed palmitoylation and prenylation signal motifs identified by our recent work with another E3-FBXL2. The indicated sequence at the top when attached to pTd-tomato vector (left) decorates plasma membranes and vesicles (right), as shown by representative confocal images. Scale bar, 10 µm.

**(E)** *FBXO10 assembly into Skip1-Cullin-1-F-box (SCF-CRL1) complex at mitochondria.* Enriched mitochondrial preparations were isolated from HEK293T cells expressing STREP-tagged FBXO10, and empty vector (EV) and processed for obtaining soluble mitochondrial lysates. Anti-STREP immunoprecipitations were performed with standard protocols. Subsequently, immunoprecipitated complexes were subject to SDS-PAGE and immunoblotting as shown. Shown is representative of three independent experiments.

**(F)** *Coomassie blue staining of samples analyzed by LFQ-mass spectrometry.* HEK293T cells expressing FLAG-tagged FBXO10, FBXO10(C953S) and empty vector (EV) were processed for obtaining enriched mitochondrial fractions by stepwise ultracentrifugation (see methods for details). A portion (~25μg) of each sample was subjected to SDS-PGAE followed by Coomassie blue staining to visualize proteins. Shown is representative Coomassie blue staining of three biological replicate samples (Mito) analyzed by LFQ-mass spectrometry. Whole cell lysate (WCL) is shown for comparison. Immunoblot of the samples with anti-FLAG, at the top, shows equal expression of FBXO10 and FBXO10(C953S) in WCLs, but enrichment of only FBXO10 in mitochondrial fractions.

**(G)** *Comparison of FBXO10 transient vs stable expression in 293T cells.* HEK293T cells were either transiently transfected with FLAG-FBXO10, FLAG-FBXO10(C953S) and empty vector (EV) plasmids-constructs or processed to obtain stable lines of indicated FLAG-tagged retro-viral constructs (see methods). Whole cell lysates were prepared from both transiently and stable expressing cells and immunoblotted as indicated.

**(H)** *Mitochondrial proteostasis via FBXO10 revealed by LFQ-mass spectrometry.* The volcano plot shows significantly altered proteins between FBXO10 and empty vector data sets (see also Fig. 1F-G). OMM proteins that showed significant protein level changes, i.e., decrease upon FBXO10, are highlighted (≥2 folds, at FDR 5%). FDR: False discovery rate. Three independent biological replicates of each FBXO10, FBXO10 (C953S) and empty vector samples were assayed by LFQ-MS/MS.

**(I)** *Mitochondrial proteostasis via FBXO10 revealed by LFQ-mass spectrometry.* Functional Annotation Clustering analysis of the significantly deregulated proteins in mitochondria expressing FBXO10 compared with FBXO10 (C953S) (FDR 5%, deregulation ≥ 2 folds) was performed using DAVID software (https://david-d.ncifcrf.gov/)[55] (data available on request). Clusters that showed enrichment score of ≥ 2 are plotted.

**(J)** *FBXO10(ΔF-box) is retained at mitochondria unlike FBXO10(C953S).* Enriched mitochondrial preparations were isolated by stepwise centrifugation from HEK293T cells expressing STREP-tagged FBXO10, FBXO10 (C953S), FBXO10(ΔF-box) and empty vector (EV). Mitochondrial preparations were processed for obtaining soluble mitochondrial lysates followed by SDS-PAGE and immunoblotting as shown.

**(K)** *$SCF^{FBXO10}$ and FBXO10(ΔF-box) are at mitochondria.* Enriched mitochondrial preparations were isolated by stepwise centrifugation from HEK293T cells expressing STREP-tagged FBXO10 and FBXO10(ΔF-box). Mitochondrial preparations were processed for obtaining soluble mitochondrial lysates. STREP-Tactin immunoprecipitations were performed with standard protocols. Subsequently, immunoprecipitated complexes were subject to SDS-PAGE and immunoblotting as shown.

**(L)** *CaaX-mutant FBXO10(C953S) and FBXO10(ΔF-box) inhibit mitochondrial membrane potential.* Measurement of mitochondrial membrane potential was carried out by flowcytometry upon TMRM treatment (100nM) in undifferentiated C2C12 cell stably expressing FBXO10, FBXO10 (C953S), FBXO10 (Δ-Fbox) and empty vector (EV). Bar graph shows quantification of mitochondrial membrane potential for the indicated samples [n=2 for EV and FBXO10 (C953S); n= 3 for FBXO10 and FBXO10 (Δ-Fbox)] p-values were calculated by student-T-test. Error bars indicate SEM.

**(M)** *FBXO10 depletion impacts mitochondrial network morphological dynamics.* FBXO10 depletion was carried out with two independent shRNAs targeted to untranslated (3'UTR) and coding (CDS) regions in C2C12 cells. Post- depletion cells were either subjected to confocal imaging or analyzed for FBXO10 mRNA depletion. Prior to imaging Mito-Tracker was added for ~30 minutes for mitochondrial network morphological analysis using confocal LSM700 microscope. Bar graph shows quantification of FBXO10 depletion by two independent shRNAs (3’UTR and CDS) analyzed by real-time RT-PCR performed as technical triplicates.

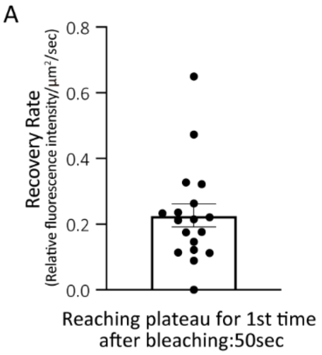


Bhat *et al.*, Figure S2

**Figure S2. FBXO10 is dynamically localized at the mitochondria.**

**(A)** *Equation for the calculation of fluorescence recovery Rate after photobleaching.* Fluorescence recovery after photobleaching (FRAP) assay for GFP-FBXO10 was carried out in Fig. 2G. The following equation was used to calculate the recovery rate.

(relative fluorescence intensity $_{\text{(reaching plateau 1st timepoint)}}$-relative fluorescence intensity$_{\text{(bleaching)}}$)/area/duration $_{\text{(reaching plateau 1st timepoint)}}$ = 0.23 0.04 (%/$\mu m^2$/sec)

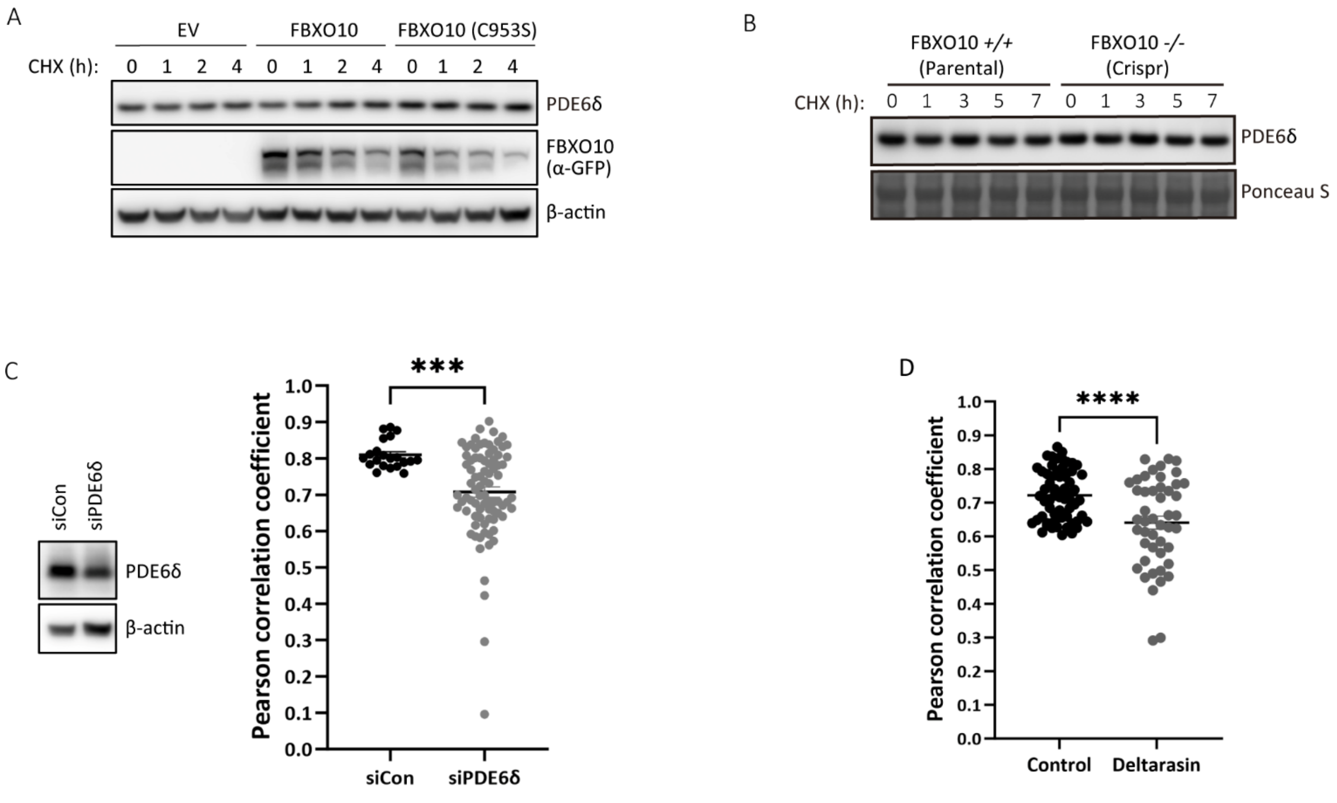


Bhat *et al.*, Figure S3

**Figure S3. PDE6δ is a cytosolic factor facilitating trafficking of geranylgeranylated-FBXO10.**

**(A)** *PDE6δ abundance is refractory to FBXO10 and FBXO10(C953S) expression.* HEK 293T expressing GFP-FBXO10, GFP-FBXO10(C953S) or empty vector (EV) were treated with cycloheximide in for indicated time points before cell harvesting. Frozen cell pellets were lysed, and whole cell lysates were processed for SDS-PAGE followed by immunoblotting as indicated.

**(B)** *FBXO10 deletion by CRISPR/CAS9 editing does not impact abundance of PDE6δ.* Parental C2C12 and CIRISPR/CAS9 deleted FBXO10 myoblast clones were differentiated for 2 days prior to treatment with cycloheximide (100 μg/ml) for the indicated time points before cell harvesting. Frozen cell pellets were lysed, and whole cell lysates were processed for SDS-PAGE followed by immunoblotting as indicated. Shown is the representative of two independent experiments.

**(C)** *PDE6δ depletion delocalizes FBXO10 from OMM.* Whole cell lysates prepared from untreated (lane 1), and siRNA treated samples shown in Fig. 3D, were processed with SDS-PAGE and immunoblotted as indicated. Scatter plots show Pearson correlation coefficients (PCC) for the colocalization of GFP-FBXO10, and mitochondria analyzed in different cells shown in Fig. 3D [ control siRNA (n=21), PDE6D siRNA (N=82)). p-values were calculated by student-T-test.

**(D)** *PDE6δ inhibition with deltarasin delocalizes FBXO10 from OMM.* Scatter plots show Pearson correlation coefficients (PCC) for the colocalization of GFP-FBXO10, and mitochondria analyzed in different cells shown in Fig. 3E [Control (n=64), deltarasin (n=47)). p-values were calculated by student-T-test.

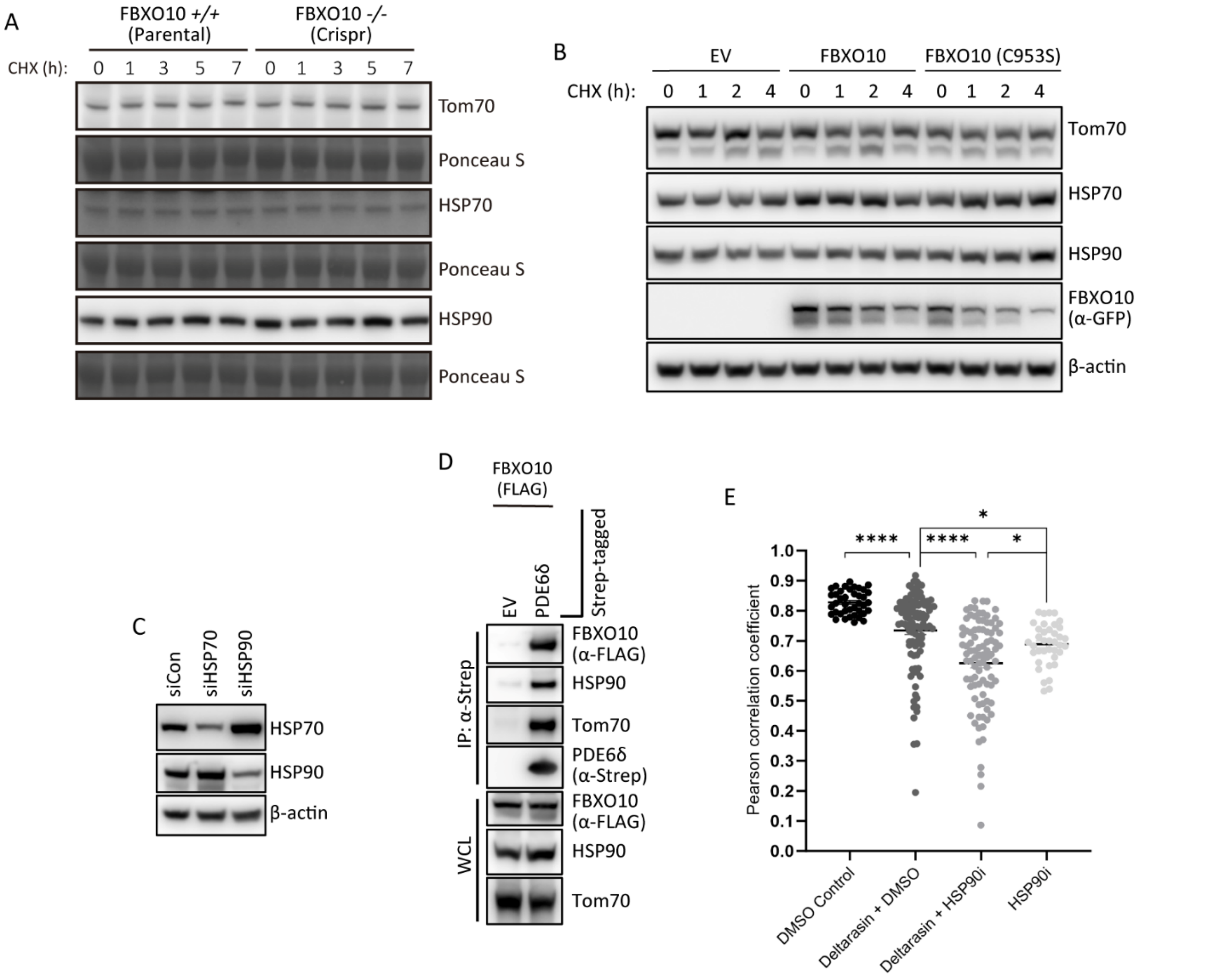


Bhat *et al.*, Figure S4

**Figure S4. HSP90 and PDE6δ ensure OMM targeting of geranylgeranylated-FBXO10.**

**(A)** *HSP90, HSP70 and TOM70 are not targets for degradation via FBXO10.* Parental C2C12 and CIRISPR/CAS9 deleted FBXO10 myoblast clones were differentiated for 2 days prior to treatment with cycloheximide (100 μg/ml) for the indicated time points before cell harvesting. Frozen cell pellets were lysed, and whole cell lysates were processed for SDS-PAGE followed by immunoblotting as indicated. Shown is the representative of two independent experiments.

**(B)** *HSP90, HSP70 and TOM70 abundance is refractory to FBXO10 and FBXO10(C953S) expression.* HEK 293T expressing GFP-FBXO10, GFP-FBXO10(C953S) or empty vector (EV) were treated with cycloheximide in for indicated time points before cell harvesting. Frozen cell pellets were lysed, and whole cell lysates were processed for SDS-PAGE followed by immunoblotting as indicated.

**(C)** *Analysis of HSP90 and HSP90 silencing.* Whole cell lysates of samples shown in Fig. 4E were immunoblotted to confirm silencing of HSP70 and HSP90. Shown is the representative of two independent experiments.

**(D)** *PDE6δ complex contains FBXO10, TOM70, HSP90 and HSP70.* STREP-tagged *PDE6δ* was co-expressed with FLAG-FBXO10 in HEK293T cells before cell harvesting. Frozen cell pellets were processed for STREP-Tactin immunoprecipitations with standard protocols. Subsequently, immunoprecipitated complexes were subject to SDS-PAGE and immunoblotting as indicated. Shown is the representative of two independent experiments.

**(E)** *Quantification of delocalization of FBXO10 away from OMM upon simultaneous inhibition of HSP90 and PDE6D.* GFP-FBXO10 expressing HeLa cells were treated with DMSO, CCT018159, deltarasin and in combination as indicated for 16 hours. Post-treatments cells were treated with Mito-Tracker for 15 minutes to decorate mitochondria before visualization by live cell confocal microscopy. Confocal images were captured using ZEIS LSM700 microscope. Pearson correlation coefficients (PCC) for the colocalization of GFP-FBXO10 and mitochondria were analyzed in different cells. Statistical analysis of plotted PPCs in scatter plots is shown (right). [n=40 for DMSO, n=38 for CCT018159 (HSP90i), n=102 deltarasin and n=90 deltarasin+ CCT018159]. p-values were calculated by student-T-test. Error bars represent SEM.

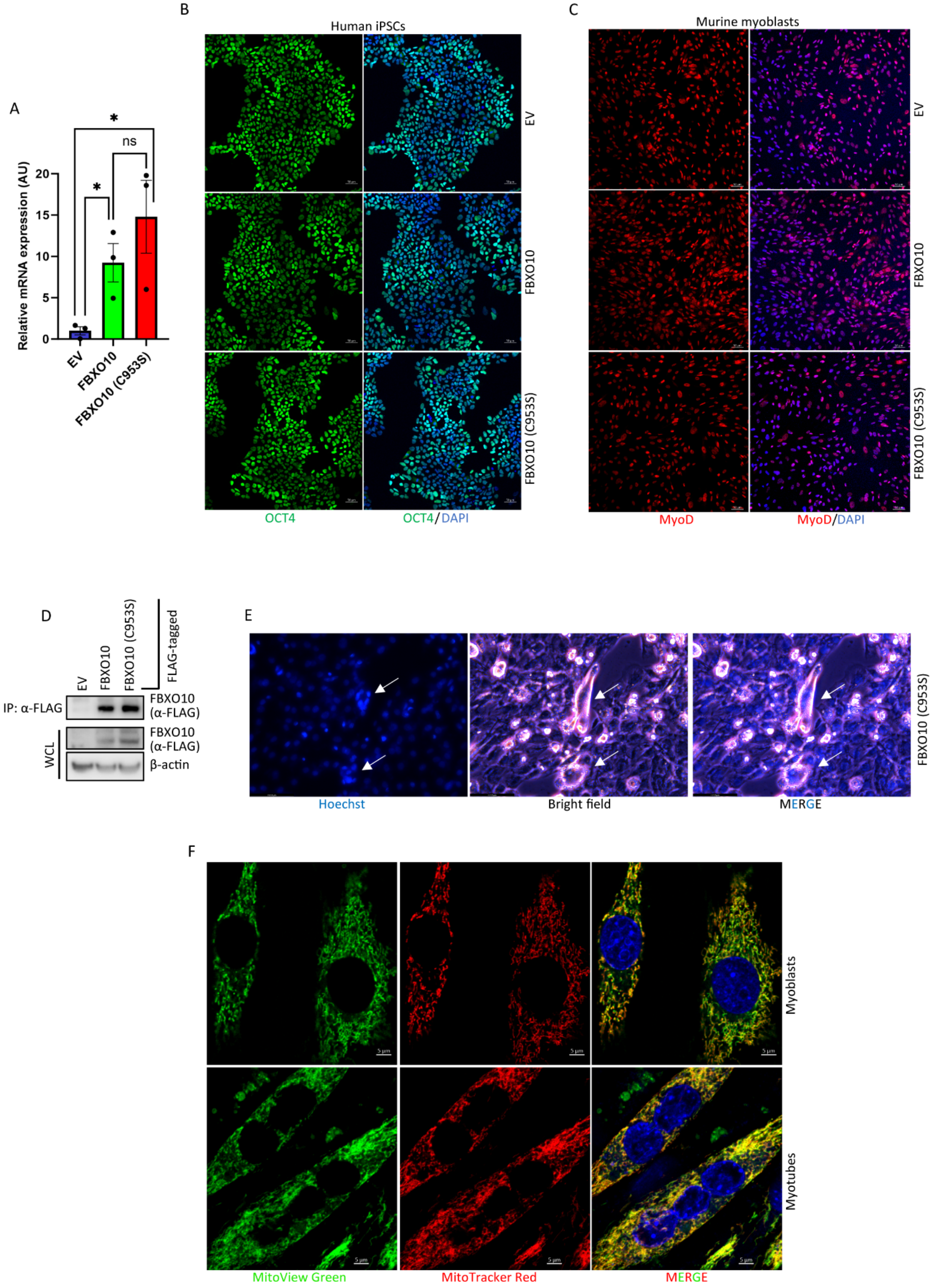


Bhat *et al.*, Figure S5

**Figure S5. Impaired mitochondria-driven myogenic differentiation in iPSCs and murine myoblasts by geranylgeranylation-deficient FBXO10(C953S).**

**(A)** *iPSCs express FBXO10 and FBXO10(C953S) stably.* Total RNA isolated from indicated human iPSCs derived samples was analyzed for FBXO10 mRNA message by RT-QPCR using specific primers (see methods section for details). Gene expression normalized to GAPDH expression ($\Delta\Delta C_q$) was calculated and plotted using GraphPad Prism software. p-values were calculated by Student-T-test. Error Bar: SEM. n=3 technical replicates.

**(B)** *Undetectable impact at progenitor stage by expression of geranylgeranylation-deficient FBXO10(C953S) in iPScs.* Human iPSCs stably expressing FLAG-FBXO10, FLAG-FBXO10(C953S) and vehicle controls (EV) were differentiated to myotubes (see protocol details in methods section). At the end point, the samples were fixed and immunostained with OCT4 antibody and Rabbit Alexa Fluor™ Plus 488 conjugated secondary antibody. DAPI was added to stain nuclei prior to visualization by ZEISS LSM700 confocal microscopy. Scale bar: 50 µm.

**(C)** *Undetectable impact to myoblast stage by expression of geranylgeranylation-deficient FBXO10(C953S) in iPScs.* Human iPSCs stably expressing FLAG-FBXO10, FLAG-FBXO10(C953S) and vehicle controls (EV) were differentiated to myoblasts (see protocol details in methods section). At the end point, the samples were fixed and immunostained with MyoD antibody and Mouse Alexa Fluor™ Plus 555 conjugated secondary antibody. DAPI was added to stain nuclei prior to visualization by ZEISS LSM700 confocal microscopy. Scale bar: 50 µm.

**(D)** *Murine myoblast C2C12 express FBXO10 and FBXO10(C953S) stably.* Murine C2C12 myoblasts stably expressing FLAG-FBXO10, FLAG-FBXO10(C953S) and vehicle controls (EV) were differentiated for 5 days for myotube formation. At the end point samples were collected for the whole cell lysate preparation followed by FLAG IP and immunoblotting against indicated proteins.

**(E)** *Geranylgeranylation-deficient FBXO10(C953S) blocks proper myotube formation.* Murine C2C12 myoblasts stably expressing FLAG-FBXO10(C953S) processed as in Fig. 5B. At the end point, the samples were treated with Hoechst, and visualized by microscopy by live cell brightfield and florescence confocal microscopy. Scale bar: 117.8 µm.

**(F)** *Mitochondrial network morphological dynamics change during differentiation.* Murine C2C12 myoblasts were differentiated for 8 days for myotube formation. At the end point samples were treated with MitoView Green, MitoTraker Red, and Hoechst to decorate mitochondria and nuclei. Representative images, captured by by ZEISS LSM700 confocal microscopy, show mitochondrial morphology changes. Shown is representative of three independent experiments. (Scale bar, 5 µm).

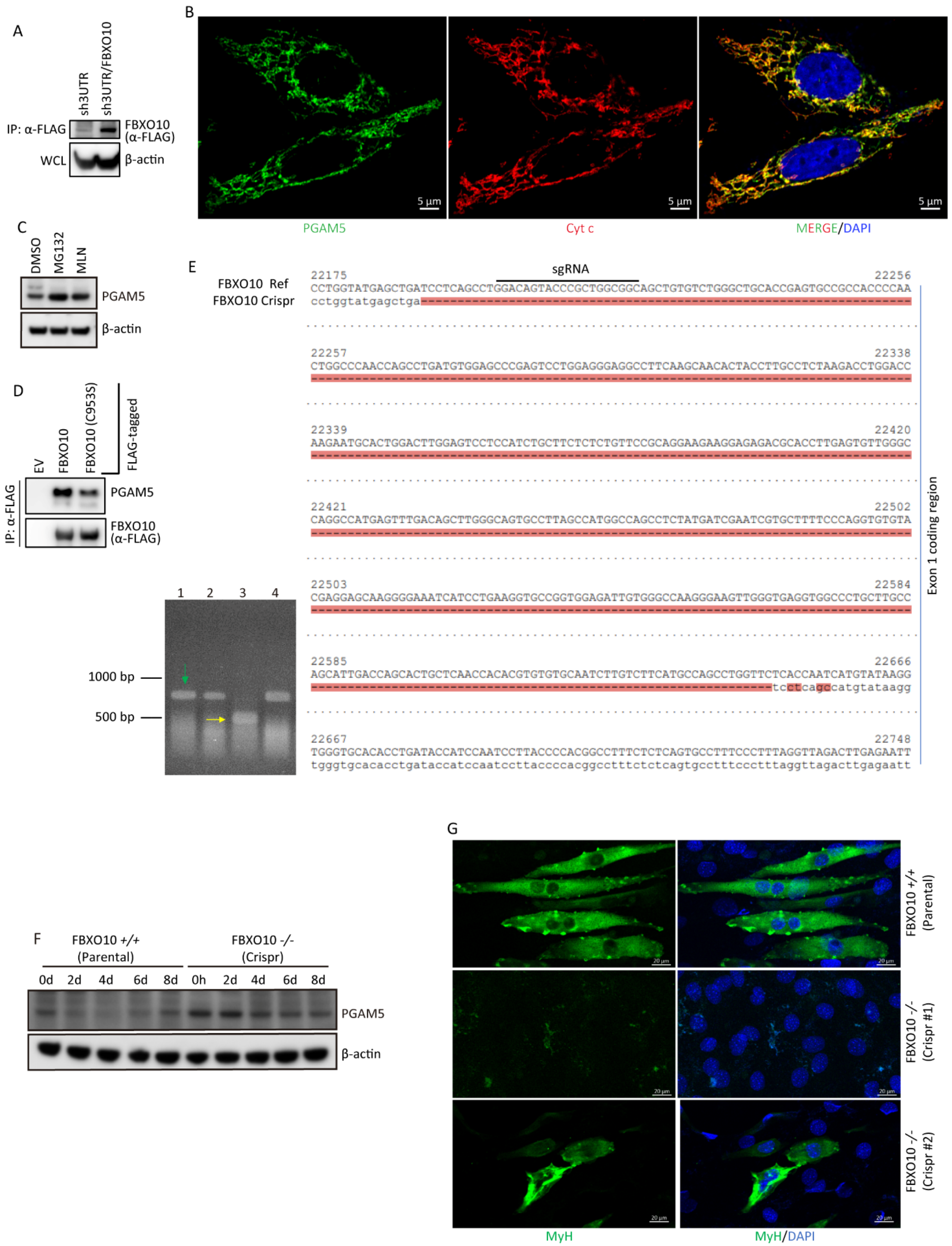


Bhat *et al.*, Figure S6

**Figure S6. FBXO10 reconstitution corrects impaired mitochondria-driven myogenic differentiation.**

**(A)** *Analysis of FBXO10 reconstitution.* Whole cell lysates were prepared from sh3'UTR and sh3'UTR/FLAG-FBXO10 myoblasts and immunoprecipitations were carried out with FLAG-M2 Agarose followed by immunoblotting to ani-FLAG antibody.

**(B)** *PGAM5 immunostaining at mitochondria.* HeLa cells were grown for ~16 hours and fixed with paraformaldehyde (4% in PBS) and permeabilized with Triton X-100 (0.1% in PBS) . Immunostaining was carried out with anti-PGAM5 and anti-Cyt c (mitochondrial marker) antibodies overnight at 4 degrees and subsequently with secondary antibodies -conjugated to Alexa-488 and Alexa-555 for 1h at room temperature. Confocal images were captured using ZEIS-LSM700 microscope. Scale bar: 5μM

**(C)** *MG312 and MLN4924 treatment increase PGAM5 levels.* C2C12 myoblasts were differentiated for 2 days. Post-2-day differentiation MG132 (10 μM) and MLN4924 (10 μM) treatments were performed for 8 hours in differentiating medium. At the end point DMSO, MG132 and MLN4924 treated samples were harvested and processed for whole cell lysate immunoblotting as indicated. Shown is the representative of the three independent experiments.

**(D)** *Loss of PGAM5 binding to FBXO10(C953S).* FLAG-tagged FBXO10, FBX010(C953S) and empty vector (EV) were expressed in HEK293T cells. Post cell harvesting anti-FLAG immunoprecipitations were performed with standard protocols. Subsequently, immunoprecipitated complexes were subject to SDS-PAGE and immunoblotting as indicated.

**(E)** *Confirmation of CRISPR-CAS9 mediated FBXO10 deletion.* (Right) FBXO10 gene was deleted with CRISPR-CAS9 editing in C2C12 myoblasts (see methods for details). FBXO10 loci around the targeted PAM site in edited and parental C2C12 cells were amplified by PCR and subjected to sanger sequencing to confirm CRISPR-CAS9 mediated FBXO10 gene disruption. (Bottom Left) PCR screening of some clones using primers around the targeted PAM site. Green arrow shows banding pattern similar to parental line whereas yellow arrow shows a major deletion in one of the clones as confirmed by sanger sequencing.

**(F)** *PGAM5 levels fluctuate during myogenic differentiation.* Myogenic differentiation in Parental C2C12 and CIRISPR/CAS9 deleted FBXO10 myoblast clone was carried out for up to 8 days before sample harvesting at indicated time points (see methods sections for details). Frozen cell pellets were lysed, and whole cell lysates were processed for SDS-PAGE followed by immunoblotting as indicated.

**(G)** *CRISPR-CAS9 mediated FBXO10 deletion blocks myogenic differentiation.* Parental C2C12 and two independent CIRISPR-CAS9 deleted FBXO10 clones (CRSPR#1 and CRSPR#2) were processed as in Fig. 6I. The samples were immunostained using A488 conjugated MyH antibody and visualized by confocal microscopy (Scale bar, 20 μm).

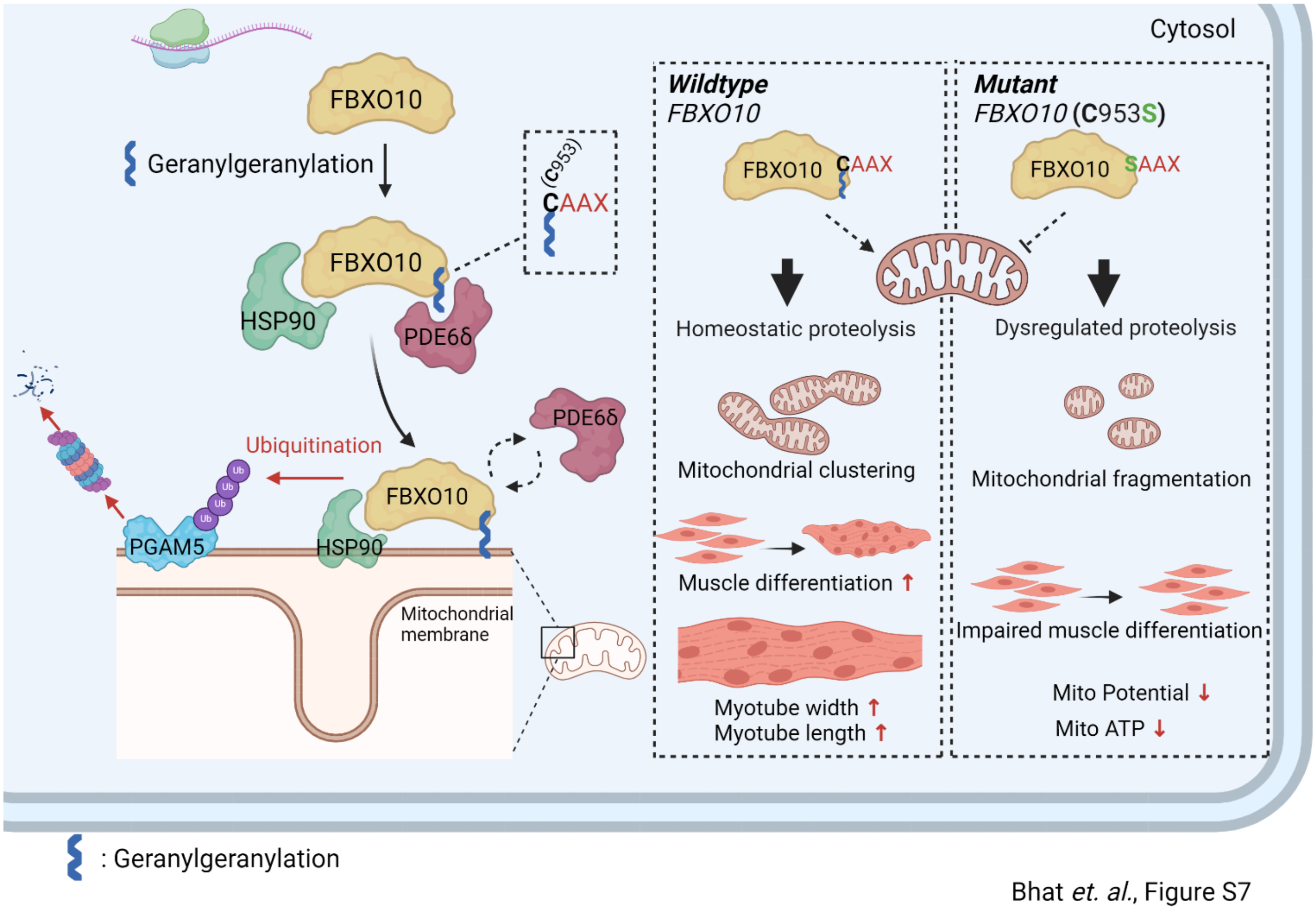


**Figure S7. Schematic: Geranylgeranylated FBXO10 is dynamically distributed at OMM to control mitochondrial proteostasis and function.**

Our new findings propose a model where in PDE6δ binds geranylgeranylated-FBXO10 to shield the isoprenoid lipid for the efficient dynamic transport in the aqueous cellular environment to and from OMM (Fig. 2, Fig. S2, Fig. 3 and Fig. S3). As FBXO10 polypeptide lacks MTS signal, HSP90 binding in coordination with PDE6δ mediated trafficking ensures specific delivery of FBXO10 to OMM *via* docking on the OMM receptor TOM70 (Fig.4 and Fig. S4). While at OMM, and depending on the cues, FBXO10 promotes selective ubiquitylation and degradation of OMM targets, for example, PGAM5 is targeted in response to myogenic differentiation cues (Fig. 6D-H and Fig. S6B, C). FBXO10 loss or re-distribution away from the mitochondria dysregulates OMM proteostasis resulting in impaired mitochondrial homeostasis which impacts myogenic differentiation (Fig.1, Fig. S1, Fig. 5, Fig. S5 Fig. 6 and Fig. S6).
